# Tools for synchronous temporal control of independent transgenes in discrete *Drosophila* tissues

**DOI:** 10.64898/2026.02.02.703225

**Authors:** Evelyne Ruchti, Brian D. McCabe

## Abstract

Exogenous binary gene regulatory expression systems are a linchpin technology critical for many model organism genetic manipulations including of *Drosophila melanogaster*. Subsequent to the initial establishment of the Gal4/UAS binary expression system, LexA/LexOp and QF/QUAS binary gene expression systems have been adopted to supplement and expand the tissue specific control of genes expression in *Drosophila*. Here, we have developed a compendium of modular vectors that enable robust, reproducible, and high throughput parallel construction of transgenes that produce either Gal4, LexA or QF2 transcription factors under specific gene enhancers and compatible partner vectors that allow transgenes to be regulated by these factors under UAS, LexOp, or QUAS control. Expanding upon this foundation, we have generated a novel hybrid binary gene regulatory system derived from QF and Gal4 – QFG4, that enables simultaneous coordinate regulation of UAS and QUAS transgenes by Gal80, allowing synchronous temporal control of independent transgene expression in distinct cells or tissues. We envision these new tools will facilitate novel and increasingly sophisticated interrogation of cell and tissue interactions particularly in contexts where temporal control is paramount.

## Introduction

The ability to reproducibly target expression of genes to defined cells or developmental time periods is an essential tool of modern model organism genetics. For example, the introduction of the *Saccharomyces cerevisiae* transcription factor, Gal4 and its recognition element the Upstream Activating Sequence (UAS) (1) into *Drosophila melanogaster* by Brand and Perrimon (2) revolutionised the speed and sophistication of genetic manipulations made possible in this model organism (1). The adaptability of this binary genetic approach – where cloned or endogenous enhancer sequences drive tissue specific expression of Gal4, which in turn activates transgenes downstream of UAS sequences – has enabled a dizzying array of investigations and applications in *Drosophila* and promoted the expansion of the binary gene regulatory approaches to other animal models (3). Generation of large libraries of tissue specific Gal4 lines (4) together with a vast range of compatible UAS transgenes has enabled genetic manipulations spanning the range from investigations of embryonic development to adult behaviour (5).

Subsequent innovations have expanded upon the original Gal4/UAS reagents to enable more precise control of transgene expression. The Gal4 protein can be broadly divided into two domains, a DNA binding domain (DBD), which recognizes and binds to the UAS sequence and an Activation domain (AD) which recruits the machinery necessary to initiate gene transcription (6). These domains are modular and separable, and can be fused with other proteins (7). For example in ‘split’ Gal4 reagents, the Gal4[DBD] and Gal4[AD] are independently produced in different patterns allowing UAS transgene expression only in tissues where the availability of both components intersects (8). The Gal4[AD] can also interact with the transcriptional repressor Gal80 which inhibits Gal4 transcriptional activity (9). This can be used to refine a pattern of UAS transgene expression if Gal80 is expressed in an intersecting pattern to Gal4 (10, 11), or to enable temporal control of Gal4 expression, essential for many studies. The latter is most commonly achieved using lines that ubiquitously express a temperature-sensitive version of Gal80 (tsGal80) (12, 13). Culturing animals at the permissive temperature (29°C) allows UAS transgene expression, while culturing animals at the restrictive temperature (18°C) inhibits transgene expression. Drug dependent modulation of Gal80 protein stability has also been developed as an alternative to temperature regulation (14–18). An important advantage of utilising Gal80 versus other approaches for binary system temporal regulation (19) is that it is compatible with existing large libraries of Gal4 lines, while other approaches require the generation of new reagents (20). The ability to control binary system regulated gene expression with accuracy in space (i.e. cells and tissues) as well as in time is highly desirable for manipulations of increasing precision and the subject of continual development (10, 15, 21–23).

A second focus of binary gene regulatory system development has been the generation of additional independent binary gene regulatory systems as a complement to Gal4/UAS. This is important to enable experiments in which transgenes (e.g. fluorescent proteins to image cells) can be expressed independently of reagents designed to simultaneously manipulate genes (e.g. RNAi constructs) either within those cells or orthogonal tissues. The second Gal4/UAS orthogonal system to be introduced in *Drosophila* was the bacteria-derived LexA/LexOp system, where a hybrid transcription factor based on LexA can induce gene expression downstream of LexOp sequences (24, 25). Transcription of LexOp and UAS regulated genes are independent of each other (24, 26). However expression levels of genes regulated by LexA/LexOp system appear less robust than Gal4/UAS, which has prompted additional modifications to generate ‘stronger’ new LexA hybrid variants (27). A second parallel expression system derived from *Neurospora crassa*, uses the QF transcription factor to promote transcription of genes downstream of QUAS sequences (28). Transcription of QUAS and UAS regulated genes are also mutually exclusive (as are QUAS and LexOp). The original QF transcription factor protein showed evidence of toxicity when expressed in some *Drosophila* tissues, prompting the generation of variants such as QF2 with reduced toxicity (27). The QF transcription factor can be inhibited by QS (analogous to inhibition of Gal4 by Gal80) which itself can be inhibited by oral administration of quinic acid (28). Both the LexA and the QF binary systems have been combined with the Gal4/UAS system in numerous studies e.g. (22, 27, 29–34) highlighting the value of multiple independent binary systems. However, a more widespread exploitation of this utility has been limited by two factors. Firstly, far fewer LexA/LexOp and QF/QUAS system lines are currently available compared to the Gal4/UAS system (35, 36). Secondly, construction of lines combining two or even three binary gene expression systems can be challenging. The location of a large number of Gal4 and LexA lines at the same genetic locus (e.g. the attP2 phiC31 landing site (37)) exacerbates this problem. Moreover, the burden of constructing lines with multiple binary systems that also allow dual temporal control, for example with both tsGal80 and QS, plus the difficulty of controlling both temperature and quinine administration in experiments, has meant that few experimental demonstrations of this dual regulatory approach exist.

To address these issues, we have generated and tested new modular vectors optimized to enable precise and robust expression of Gal4, LexA and QF2, in addition to compatible UAS, LexOp and QUAS vectors, all of which are phiC31 landing site compatible (37). We showed that these lines can be used to produce similar cell type specific patterns of Gal4, LexA and QF2 expression, with the same enhancer element, at different genomic locations. We also generate a new hybrid binary gene expression system, derived from QF and Gal4, which allows simultaneous regulation of two binary gene expression systems by a single Gal80 element, while maintaining distinct tissue expression. We envision these new tools will facilitate novel types of studies that employ the coordinated temporal and spatial control of multiple genes in distinct patterns of expression to probe inter-cellular and inter-tissue interactions throughout animal development and adulthood.

## Results

### pBID2 enhances expression of transgenes

We previously generated a modular series of vectors (attB Insulated *Drosophila* - pBID) that enabled reproducible expression of transgenes from diverse attP insertion sites (38). This was enabled by the introduction of gypsy insulator sequences (39) surrounding the insertion site of the vector, coupled with the employment of a *Drosophila* Synthetic Core Promoter (DSCP) minimal promoter (40). This was designed to avoid unwanted Gal4-independent expression of the Gal4 which has previously been associated with the hsp70 minimal promoter due to it’s interaction with insulator sequences (41). However, a disadvantage of this vector series is that the DSCP minimal promoter generally induces lower levels of transgene expression than the hsp70 minimal promoter (41). While this can have advantages for some experiments (42), in other contexts lower levels of transgene expression can limit utility. To address this limitation we sought to generate an improved vector series, pBID2, with enhanced expression levels. Like the original pBID vectors (pBID1), the pBID2 vector series maintains mini-white transgenic animal selection. All pBID2 vectors are compatible with phiC31 genomic integration (43) and include Gateway (44) or ‘destination’ cassettes to enable high throughput parallel cloning (Figure 1D). To accommodate alternative cloning strategies, a similar pBID2 series compatible with conventional restriction enzyme cloning via a multiple cloning site was also generated. To increase expression of pBID2 relative to pBID1, we took advantage of a number of previously described innovations to increase the expression of *Drosophila* transgenes. We tested these modifications in a novel UAS vector, pBID2_UAS. In pBID2_UAS, we increased the number of UAS binding sequences to 20 copies (41), replaced the 3’ Untranslated Region (3’UTR) derived from hsp70 with a p10 3’UTR (45) and added a Syn21 translational enhancer sequence immediately before the start codon of the transgene (45) (Figure1A). To evaluate the impact of these modifications, we compared the levels of destabilised GFP (dsGFP) (46) produced from the original pBID1_UAS vector and the modified pBID2_UAS vector. Both constructs were integrated into identical phiC31 attP sites (attP40 (47)) and expressed using the same Gal4 driver line (GMR-Gal4 (48)), and protein levels assessed by immunoblotting. dsGFP has *∼* 2 hour half-life, hence serving as a sensitive reporter for levels of transgene expression (46). We observed a 155 % (p < 0.01) increase in the amount of dsGFP expressed from pBID2_UAS compared to dsGFP produced from pBID1_UAS (Wang et al., 2012) (Figure 1B-C). Thus as desired, pBID2_UAS allows for significantly enhanced transgene expression levels compared to the previous pBID1 vector while still retaining the advantage of transgene insulation, ensuring more consistent genomic site-to-site expression and avoiding unwanted independent Gal4 ‘leak’ (14, 41, 47, 49).

**Figure 1.**
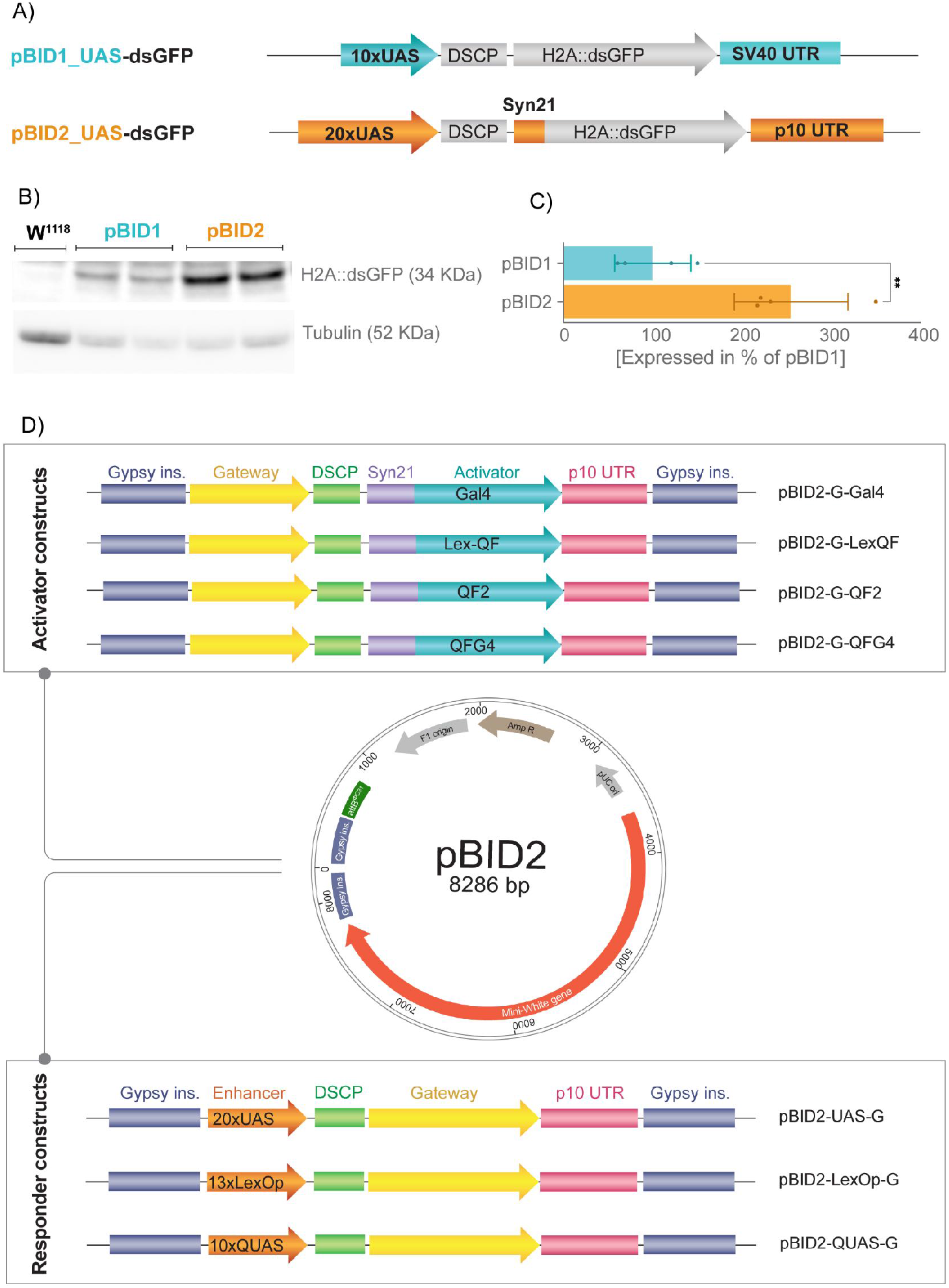
pBID2 vector design. A) Schematic of the pBID1 vector design compared with the pBID2 vector. B) Expression of destabilized GFP protein in adult heads from pBID1_10xUAS and pBID2_20xUAS inserted on chromosome II (attP40) and driven by GMR-Gal4 (*n*=4). The loading control is *α*-Tubulin. C) Quantification expression level, unpaired T-test. **p value: 0.0068. D) Overview of the complete pBID2 vector collection. Center: Map of the pBID2 vector [to scale]. pBID2 includes a mini-white gene, a *ϕ*C31 integrase compatible attB sequence (attB*ϕ*C31), an ampicillin resistance (AmpR) gene and gypsy insulator (Gypsy Ins.) sequences flanking insertion site. Top: Schematic of activator cassettes. [not to scale]. A Gateway attR destination cloning cassette (for insertion of gene regulatory sequences), is followed by a DSCP synthetic core promoter, a Syn21 translation enhancer, the activator binary transcription factor, followed by a p10 3’ untranslated region (UTR). Bottom: Schematic of responder cassettes [not to scale]. Multiple copies of the transcription factor binding sequence are followed by a DSCP synthetic core promoter, a Gateway attR destination cloning cassette (for insertion of the transgene), followed by a p10 3’UTR.

### pBID2 vector for LexOp and QUAS controlled gene expression

Building upon the foundation of the pBID2 backbone and in order to expand our vector set for additional binary gene expression systems, we next sought to generate analogous vectors with LexOp or QUAS binding sites to enable transgene control by LexA or QF respectively. We introduced 13xLexOp binding sites (45) or 10xQUAS binding sites (28) into the pBID2 backbone to generate pBID2_LexOp and pBID2_QUAS respectively (Figure 1D). To evaluate these vectors, we generated three novel nucleus-membrane markers. The first was a H2A::RFP-T2A-TFP::CAAX construct, where TagRFP-T (50, 51) was fused to the nuclear Histone 2A protein (H2A) and the Teal Fluorescent Protein (TFP) (52) was fused to the CAAX membrane-targeting motif (45, 53, 54). This marker was cloned this into pBID2_LexOp and when expressed under the control of the vGlut-LexA line (21), resulting in strong nuclear RFP and membrane localized TFP expression in glutamatergic neurons as expected (Figure 2A). Next, we also generated an H2A::GFP-T2A-mKO2::CAAX construct where sfGFP (55) was fused to the H2A protein and the Kusubira orange fluorescent protein (mKO2) (56) was fused to the CAAX motif. This marker was cloned into pBID2_QUAS and pBID2_UAS. When expressed under the control of the subset neuronal driver HB9-QF2 (57) and HB9-Gal4 (58) respectively, we observed expression in the expected neuronal pattern for both markers (Figure 2B-C). A key advantage of this approach is that all three of these pBID2 vectors employ gateway cloning cassettes (44, 59) and hence, UAS, LexOp and QUAS transgenes can be generated simply and in parallel from the same entry clone (44) aiding the rapid expansion of comparable *Drosophila* lines.

**Figure 2.**
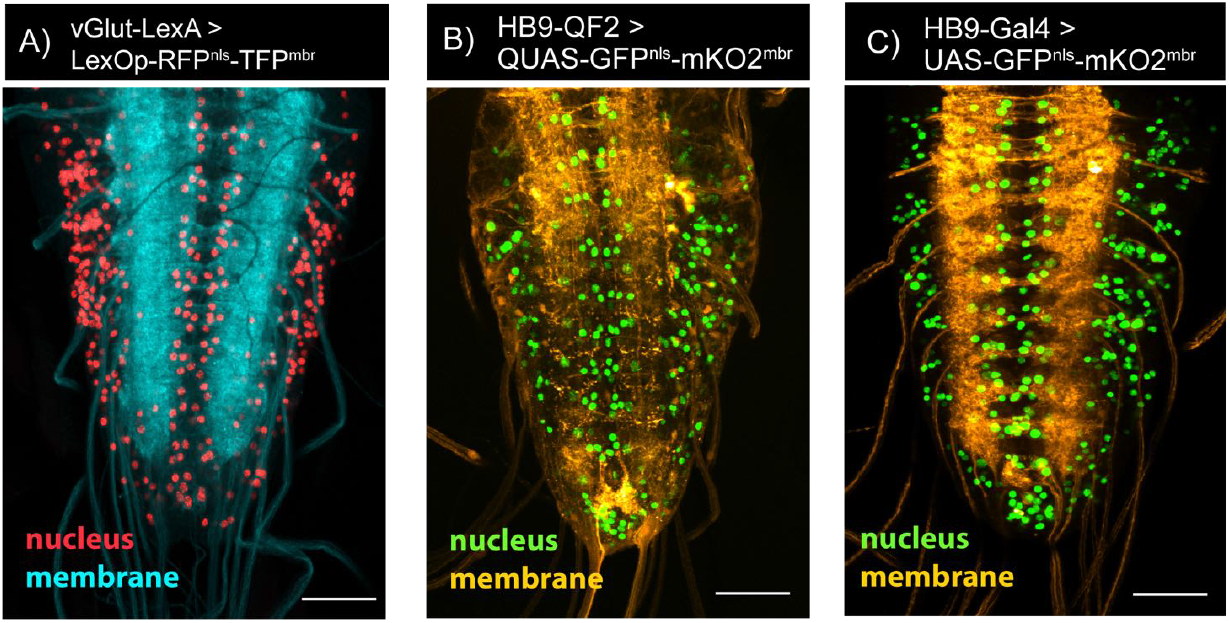
pBID2 responder construct. *Drosophila* L3 larva VNC expression of responder constructs made for three binary systems in pBID2 vector. A) vGlut-LexA driving 13xLexOp-H2A::RFP-TFP::CAAX reporter glutamatergic neurons. Nuclei appear in red and membranes in blue. B) HB9-QF2 driving 10xQUAS-H2A::GFP-mKO2::CAAX in a subset of motor neurons. C) HB9-Gal4 driving 20xUAS-H2A::GFP-mKO2::CAAX reporter in a subset of motor neurons. C, D) Nuclei appear in green and membranes in orange. Scale bars: 50 µM.

### T2A but not P2A allows efficient multiple peptide production in *Drosophila* neurons

Two ribosomal skipping sequences have been employed in *Drosophila*: T2A derived from Thosea Asigna virus (60) and P2A derived from Porcine Teschovirus (61). Both sequences induce ribosomal skipping enabling two or more peptides to be translated from a single mRNA sequence (62). As mentioned above, we used ribosomal skipping sequences (63–65), such as T2A, to enable the production of multiple independent peptides from a UAS, LexOp or QUAS single construct. We have recently shown that nucleus targeted fluorescent reporters enable the robust and accurate quantification of cell numbers in whole tissues such as the central nervous system (CNS) (66) while membrane reporters are also widely used to label cell processes including those of neurons (67). The production of multiple peptides from a single open reading frame has many uses for cell specific manipulations with the facility of a generating a single transgenic construct (68, 69) and has been used both for transgenes (70) and transgenic insertions into endogenous loci (21) (Figure 3A). To compare the efficacy of these ribosomal skipping sequences in *Drosophila* neurons, we generated two nucleus-membrane marker constructs designed to simultaneously express H2A::GFP and mKO2::CAAX (56). One of these sequences was separated by a T2A sequence [pBID2_UAS-H2A::GFP-T2A-mKO2::CAAX] and the other was separated by a P2A sequence [pBID2_UAS-H2A::GFP-P2A-mKO2::CAAX]. The two transgenes were incorporated in the same landing site [attP JK66B] (71). We then expressed these constructs with HB9-Gal4 (58). We found that in animals expressing H2A::GFP-**T2A**-mKO2::CAAX, GFP was localised to neuron nuclei and mKO2 was found at neuronal membranes as expected, with little to no overlap between these fusion proteins (Figure 3B). In contrast, when we examined animals expressing H2A::GFP-**P2A**-mKO2::CAAX, we found significant levels of mKO2 localised to the nucleus in addition to the neuronal membrane (Figure 3C). This suggested that the ribosomal skipping process was not efficient in the P2A construct resulting in the translation of a fusion protein of H2A::GFP and mKO2::CAAX that aberrantly localised to the nucleus. To determine if this was indeed the case, we examined the peptides produced by each construct in the larval CNS, using protein blotting against GFP. We found that the UAS-H2A::GFP-**T2A**-mKO2::CAAX transgene produced a peptide with the expected molecular weight for H2A::GFP. In contrast, the UAS-H2A::GFP-**P2A**-mKO2::CAAX construct produced 2 peptides, one with the expected molecular weight for H2A::GFP, and a second much larger peptide with a molecular weight consistent with the H2A::GFP-mKO2::CAAX read-through fusion protein (Figure 3D). Therefore, both our fluorescence protein localisation results and protein blotting were consistent with inefficient ribosomal skipping by the P2A sequence in stark contrast to the apparently completely efficient T2A sequence. Our results are consistent with observations from Human, Mouse and *Drosophila* cell lines reporting context-specific efficacy of ribosomal skipping sequences (68, 70, 72). We conclude that in *Drosophila* neurons, T2A provides more reliable ribosomal skipping than P2A and is therefore the preferred sequence for constructs designed to express multiple independent peptides in this cell type.

**Figure 3.**
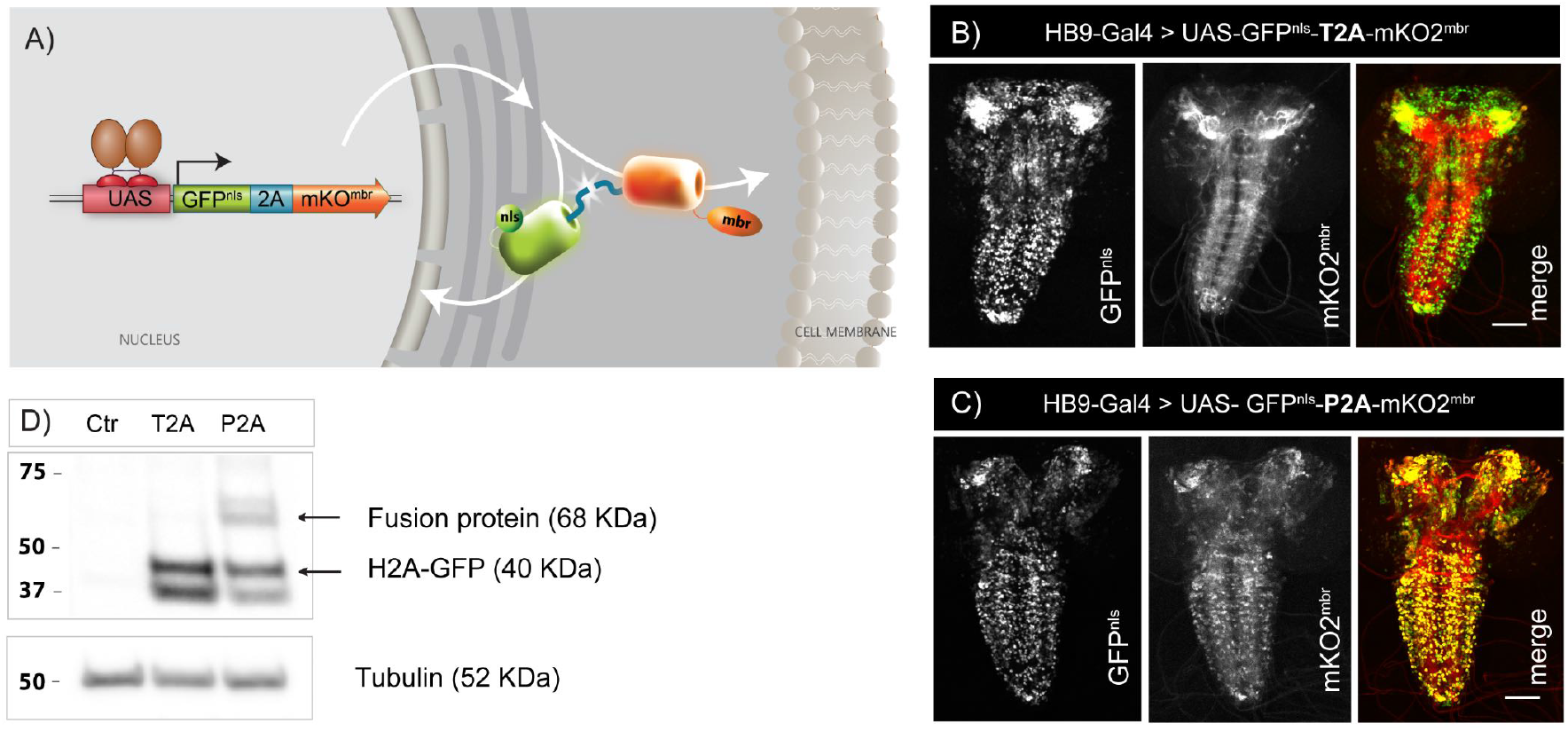
Comparison between P2A and T2A ribosomal skipping sequence. A) Schematic of nucleus-membrane marker transgene. Expression of transgene upon the binding of a binary driver (e.g. Gal4) to its corresponding regulator sequence. A single mRNA is synthesised and split at the ribosomal skipping sequence (represented in blue), giving rise to two independent reporter proteins. The reporter tagged with H2A is localised to the nucleus while the CAAX tagged reporter is localised to the membrane. B) *Drosophila* L3 larva VNC showing the distribution of GFP nucleus tagged reporter and mKO2 membrane tagged reporter after cleavage at T2A or P2A sites. Scale bar: 50 µM. C) Representative western blot of adult fly heads extract showing H2A::GFP protein after cleavage at T2A or P2A site. 68 KDa band indicates undesired fusion protein.

### pBID2 vectors for enhancer control of Gal4, LexQF and QF2 expression

We next sought to leverage the pBID2 backbone to produce modular vectors enabling the expression of Gal4, LexA or QF transcription factors under the regulation of gene enhancer sequences. To do this, we introduced either Gal4 (variant Gal4.2) (21), the LexA hybrid LexQF (27) or the QF variant QF2 (27) into the pBID2 backbone preceded by the DSCP minimal promoter and a Gateway destination cloning cassette (or alternately a Multiple cloning site (MCS) (Figure S2) to produce pBID2_Gal4, pBID2_LexQF and pBID2_QF2 respectively. To evaluate these vectors, we sought to identify a novel enhancer sequence that would label a discrete cell type to enable easy visualisation and comparison. Larval motor neurons and their associated neuromuscular junction (NMJ) synaptic terminals have been investigated in a large number of *Drosophila* neuroscience and neurological disease studies (73, 74). However, while several enhancer trap lines have been identified that label motor neurons (75, 76), to our knowledge, no transferable enhancer sequence that specifically labels all of these neurons has been isolated. *Drosophila* motor neurons are glutamatergic and enhancer regions of the *vGlut* gene that encodes the vesicular glutamate transporter (vGlut) can label all motor neurons in addition to other glutamatergic central neurons (77). Intriguingly some portions of the vGlut enhancer have been reported to express in subsets of adult leg motor neurons (78). Upon examining the vGlut locus, we cloned a putative enhancer region of the vGlut regulatory sequence [Chromosome 2L:2,403,206 to 2,403,845 [+]] (Figure 4A) and inserted it into pBID2_Gal4, pBID2_LexQF and pBID2_QF2. We generated transgenes with all three vectors and subsequently examined their expression pattern. Using these lines to express UAS, LexOp or QUAS reporter constructs, we observed expression in all motor neurons in abdominal segments of third instar larvae (Figure 4B) as evidenced by labelling of abdominal neuromuscular junction terminals (Figure 4C). To verify if this labelling was exclusively glutamatergic, we expressed a fluorescent reporter in VGMN expressing neurons and co-stained the labelled terminals with an anti-vGlut antibody (S3D). As expected, all labelled terminals were all glutamatergic. We therefore dubbed this enhancer VGMN (**VG**lut **M**otor **N**euron).

**Figure 4.**
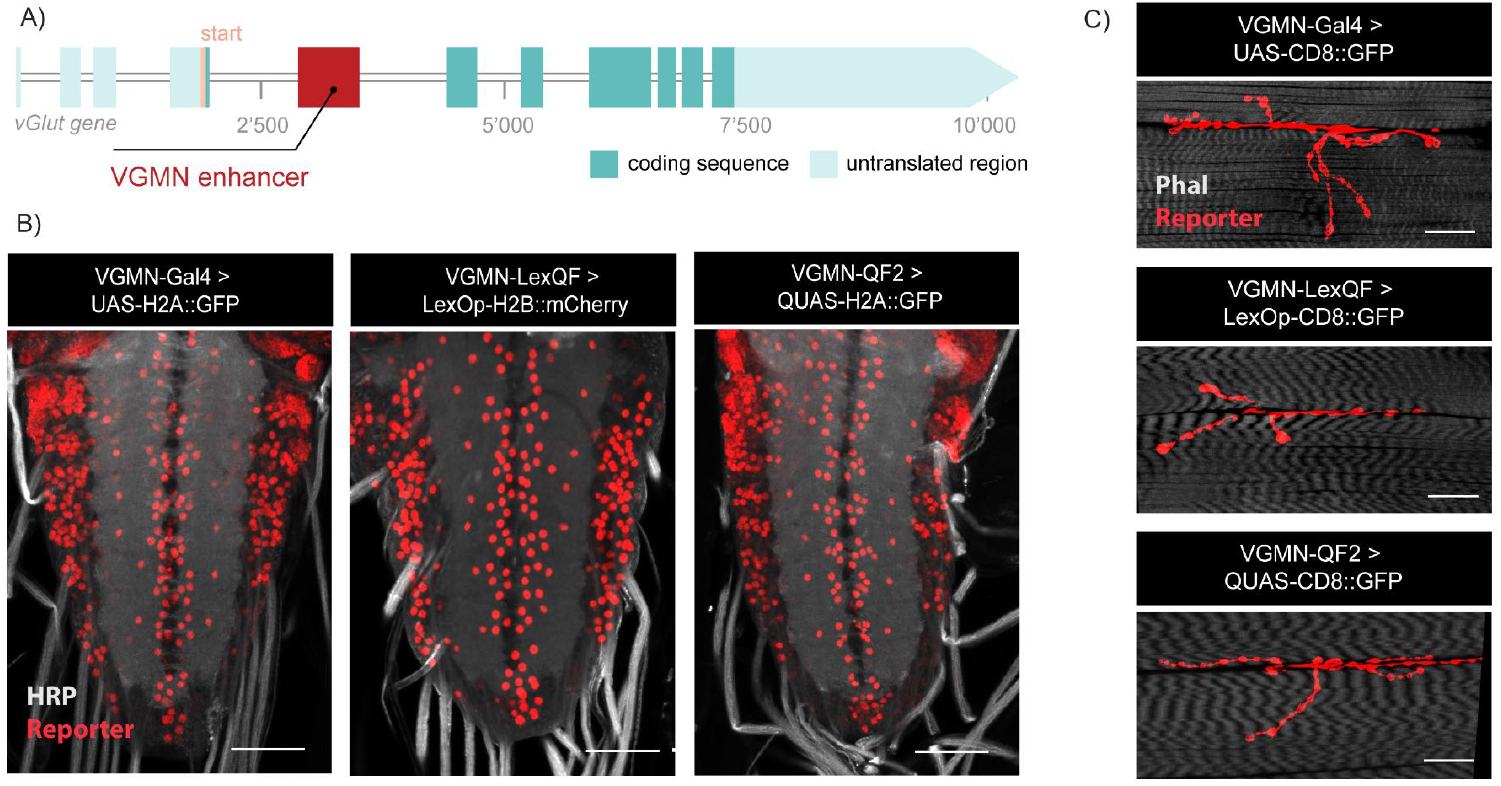
VGMN regulator inserted in pBID2 activator vector. A) Schematic showing the localization of VGMN regulator sequence (in red) in vGlut gene intronic region. B) *Drosophila* L3 larva VNC showing nuclear fluorescent reporter expression under the control of VGMN driver for the 3 binary system [VGMN-Gal4, UAS-H2A::GFP], [VGMN-LexQF, LexOp-H2B::mCherry] and [VGMN-QF2, QUAS-H2A::GFP]. Scale bar: 50 µM. C) *Drosophila* L3 larva motor neurons terminals showing membrane fluorescent reporter expression under the control of VGMN drivers for the 3 binary system [VGMN-Gal4, UAS-CD8:GFP], [VGMN-LexQF, LexOp-CD8::GFP] and [VGMN-QF2, QUAS-CD8::GFP]. Scale bar: 25 µM.

Next, to determine if the VGMN enhancer also labelled glutamatergic interneurons in the ventral nerve cord (VNC), we compared the number of neurons labelled by the pan-glutamatergic driver, OK371-Gal4 (77), to the number labelled by VGMN-LexQF within A4 to A6 segments using a nucleus reporter (66). We found that VGMN-LexQF labelled 333 +/- 23 neurons while OK371-Gal4 labelled 385 +/- 25 neurons in these segments. (Figure S3A-C) Therefore, the VGMN vGlut subset enhancer labels all motor neurons but not all glutamatergic neurons, at least in abdominal segments of larva. In contrast to the existing widely used reagent, OK371-Gal4. Taken together, pBID2 Gal4, LexQF and QF2 vectors, analogous to pBID2 UAS, LexOp and QUAS vectors, enable the rapid generation of new lines with reproducible expression patterns for multiple binary systems in parallel.

### QFG4, a novel QF hybrid regulatable by Gal80

The QF/QUAS binary gene expression system can be inhibited by QS (28), in contrast to the LexA/LexOp binary system for which no such analogous negative regulator is currently employed (24). However, due to the additional burden to construct lines with Gal80 and QS, plus the sparsity of tissue specific QS lines (in contrast to Gal80 (10)) simultaneous dual negative regulation of both systems is rarely utilised. To address this limitation, we sought to generate a hybrid QF variant that could be also regulated by Gal80 in tandem with Gal4 to exploit both existing Gal80 libraries and Gal80 temporal control tools (10, 14–16, 18). To do this, we fused the DBD of QF (amino acids 1-183) (79) to the AD of Gal4 (amino acids 148 - 882) (6), the region of Gal4 that enables negative regulation by Gal80 (80). We dubbed this QF[DBD]::Gal4[AD] hybrid, QFG4. We introduced QFG4 into a pBID2 vector to generate pBID2_QFG4 and generated lines where QFG4 was expressed under the control of the VGMN enhancer. We first examined if these lines were capable of inducing QUAS regulated transgenes. Similar to VGMN-QF2 lines, VGMN-QFG4 lines also showed specific and strong expression of QUAS-H2A::GFP in larval motor neurons (Figure 5A). In contrast, VGMN-QFG4 displayed no induction of UAS-H2A::GFP unlike VGMN-Gal4 (Figure 5B) which promoted strong UAS-H2A::GFP expression. Thus, as expected, QFG4 specifically regulates the expression of QUAS but not UAS transgenes. Given this property and to further extend the QFG4 toolkit, we also generated a pan-neuronal Synaptobrevin (nSyb) -QFG4 line by cloning this enhancer (40) into pBID2. Using a QUAS nuclear reporter, we confirmed that this line also reproduces the labelling pattern previously observed for the other nSyb driver lines (Figure S4).

**Figure 5.**
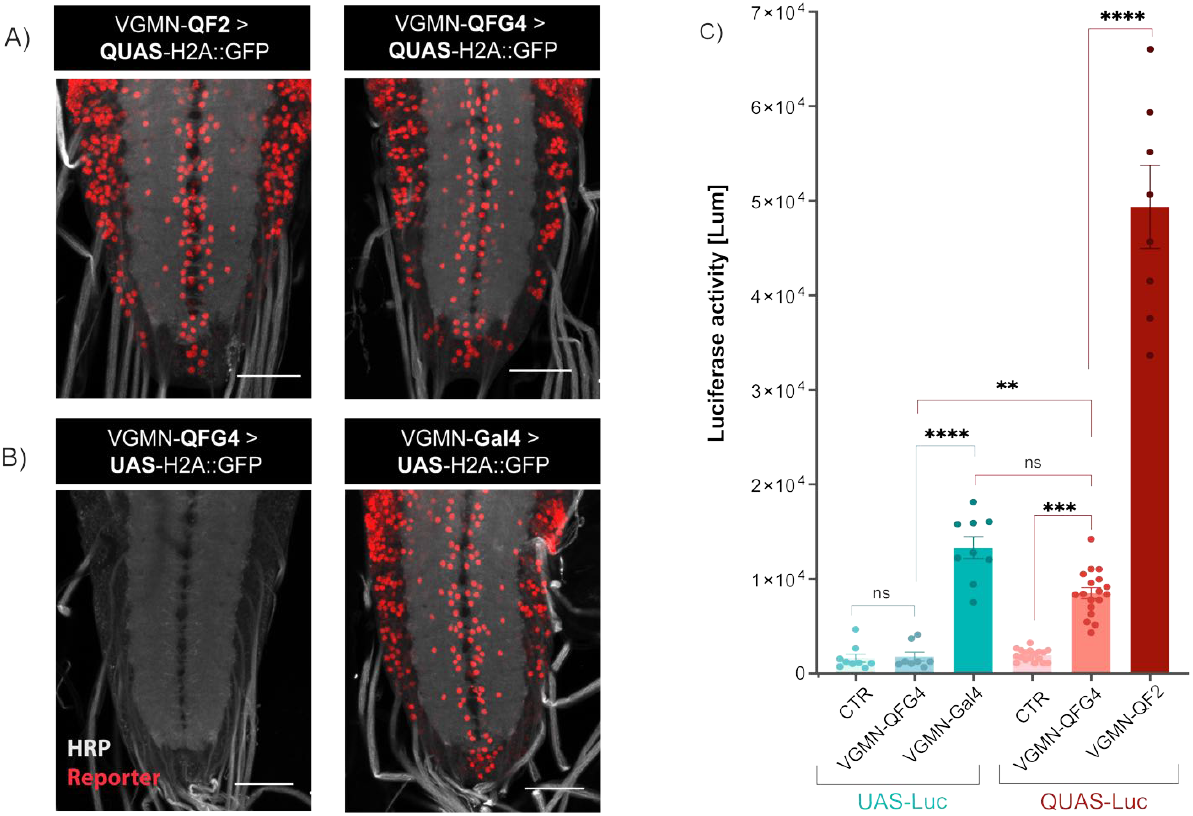
pBID2 vector with a chimeric QFG4 activator. *Drosophila* L3 larval VNC confocal images showing motor neuron expression from: A) VGMN-QF2 and VGMN-QFG4 driving QUAS-H2A::GFP and B) VGMN-QFG4 and VGMN-Gal4 driving UAS-H2A::GFP. Scale bar: 50µM. C) Expression of VGMN-QF2, VGMN-QFG4, VGMN-Gal4 compared to control (w1118) are detected by luciferase assay in *Drosophila* adult head extracts. Measured lumens from each extract are expressed as absolute values and are normalized to their corresponding protein concentration. n=8 from 3 individual experiments. Statistical analysis, One-Way Anova, Tukey’s multiple comparison test. ****p0.0001; n.s. p>0.05.

While the pattern of QF2 and QFG4 expression under the VGMN enhancer looked similar, we wondered if the transgene expression levels were comparable. To determine this, we compared, using QUAS and UAS luciferase transgenes (2, 28), the level of transgene expression induced by VGMN-QFG4, VGMN-QF2 and VGMN-Gal4 by quantitation of luciferase activity from dissected CNS tissues. Consistent with what was observed with immunostaining, VGMN-QFG4 specifically activated the QUAS but not the UAS luciferase reporter. We found that luciferase activity was 3.7 fold (p<0.001) higher when expressed using VGMN-QF2 compared to VGMN-Gal4 (Figure 5C). In contrast, luciferase activity was not significantly different when expressed from VGMN-QFG4 compared to VGMN-Gal4 (Figure 5C) and dramatically lower compared to VGMN-QF2. This large difference in expression could perhaps explain, in addition to other factors (22), some of the residual toxicity still observed in the use of some QF2 lines as QF2/QUAS transgene expression levels are much higher compared to similar Gal4/UAS experiments.

### QFG4 can be negatively regulated by Gal80

Next, we sought to determine if QFG4 could be negatively regulated by Gal80. To examine this, we employed an Anion-Gated-Channel Rhodopsin fused to YFP (GtACR1::YFP) under the control of UAS or QUAS sequences, which acts as a potent inhibitor of neuronal activity when expressed (81) (Figure 6A). To enable the visualisation of transgene expression, we expressed UAS or QUAS regulated GtACR1::YFP under the control of VGMN-Gal4 or VGMN-QFG4 respectively, with or without ubiquitously expressed Tubulin-Gal80 (10). Confocal imaging revealed no expression of UAS-GtACR1::YFP induced by the VGMN-Gal4 in the presence of Gal80, while controls without Gal80 showed robust expression (Figure 6B). We then examined QUAS-GtACR1::YFP expression in the presence of our novel QFG4 construct. Similar to VGMN-Gal4 regulated expression, we also saw robust expression of QUAS-GtACR1::YFP by VGMN-QFG4 in the absence of Gal80, yet extremely low expression of QUAS-GtACR1::YFP by VGMN-QFG4 when Gal80 was present. Next, we examined if GtACR1::YFP activity was also inhibited when Gal80 was expressed. When UAS-GtACR1::YFP was expressed by VGMN-Gal4 in the absence of Gal80, illumination of motor neurons with 515 nm wavelength light inhibited neuronal activity as expected (Figure 6C). This inhibition was completely abolished when Gal80 was present. Similarly, when QUAS-GtACR1::YFP was expressed by VGMN-QFG4 in the absence of Gal80, illumination of motor neurons also potently inhibited neuronal activity. However neuronal activity was unaffected by illumination when Gal80 was co-expressed together with VGMN-QFG4 and QUAS-GtACR1::YFP (Figure 6C). Thus, similarly to Gal4, we show that QFG4 is effectively inhibited by Gal80.

**Figure 6.**
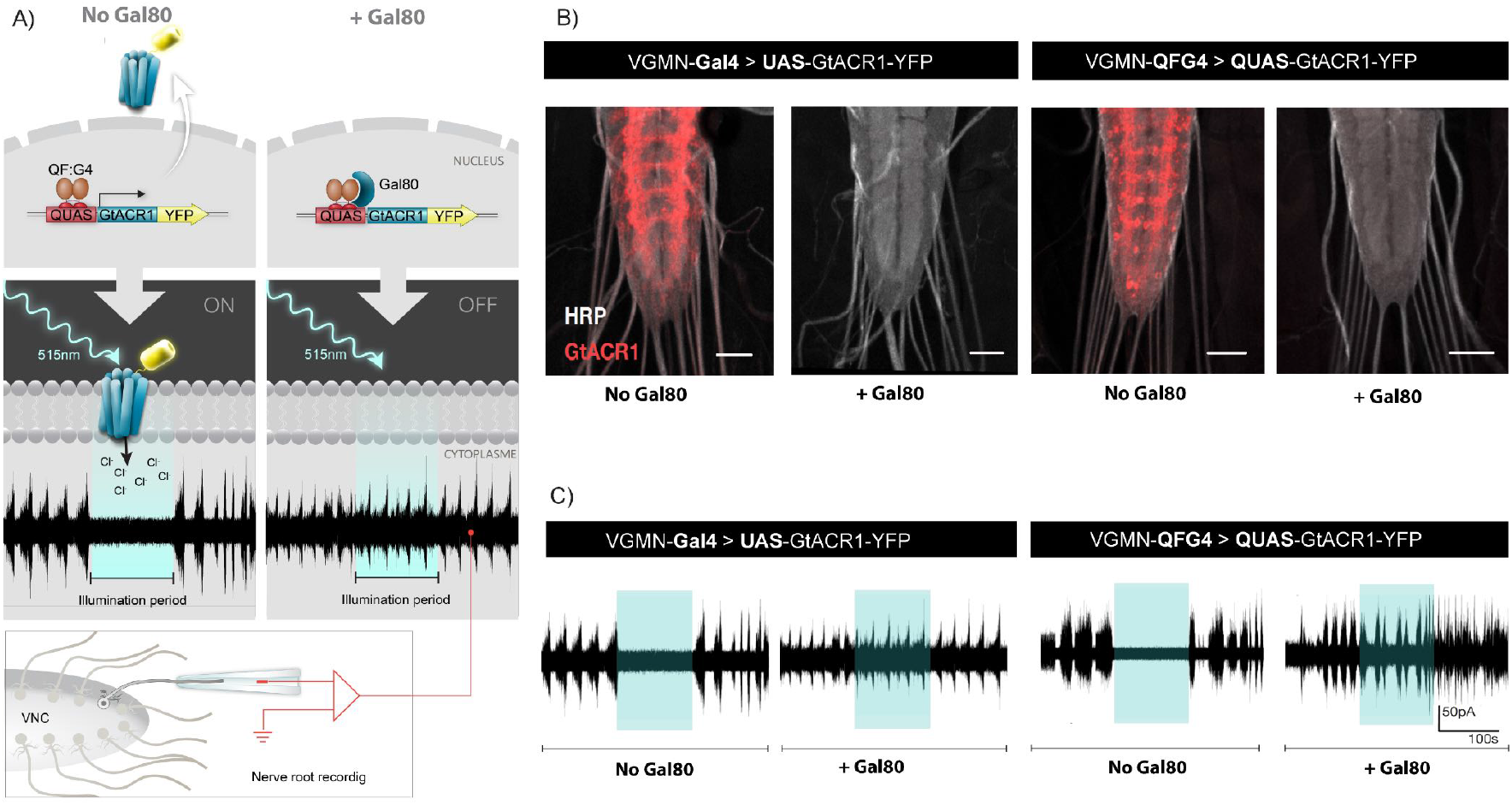
QFG4 can be negatively regulated by Gal80. A) Left panel: Schematic of QFG4 binding QUAS-GtACR1, in absence of Gal80. GtACR1 is a membrane-expressed protein that inhibits neuronal activity by mediating chloride influx upon illumination at 515 nm. Rights panel: Schematic of QFG4 binding QUAS-GtACR1, in absence of Gal80. B) Left: Motor neurons expressing GtACR1 (red) under the control of VGMN-Gal4, in absence and in presence of Gal80. Right: Motor neurons expressing GtACR1 (red) under the control of VGMN-QFG4, in absence and in presence of Gal80. Scale bar: 50 µM. C) Representatives traces of nerve roots spontaneous activity recorded from VNC motor neurons in panel B.

### Gal4 and QFG4 can be simultaneously regulated by temperature sensitive Gal80

Given the robust inhibition of QFG4 by Gal80, we next asked whether QFG4 dependent transcription could be simultaneously regulated alongside Gal4 in distinct tissues by temperature-sensitive Gal80 (tsGal80), a widely used temporal control approach for Gal4 expression (82) (Figure 7A). To address this question, we generated lines that expressed QUAS-mCherry6x (83) in motor neurons under VGMN-QFG4 control, UAS-CD8:GFP (Lee and Luo, 1999) in muscles under MHC-Gal4 control (84), and that also ubiquitously produce tsGal80 (82). As a preliminary test, we cultured animals in parallel, at the tsGal80 permissive temperature of 29°C and independently at the tsGal80 restrictive temperature of 18°C. L3 larva revealed complete absence of both UAS and QUAS reporters when raised at 18°C, while strong expression of both reporters could be observed when raised at 29°C. (Figure S5). We then repeated the experience on adult flies, where animals were raised at 18°C and subsequently moved to 29°C after 7 days. (Figure 7B) When cultured at the tsGal80 restrictive temperature of 18°C, we observed no expression of the UAS-reporter and very low expression of the QUAS-reporter. In contrast, when we transferred these animals to the tsGal80 permissive temperature of 29°C, we observed robust expression of the QUAS mCherry reporter in motor neurons and the UAS GFP reporter in muscles as expected (Figure 7B). Thus, lines expressing QFG4 together with Gal4 under tissue specific control enable coordinate regulation of both QUAS and UAS transgenes in distinct tissues by a single tsGal80 element.

**Figure 7.**
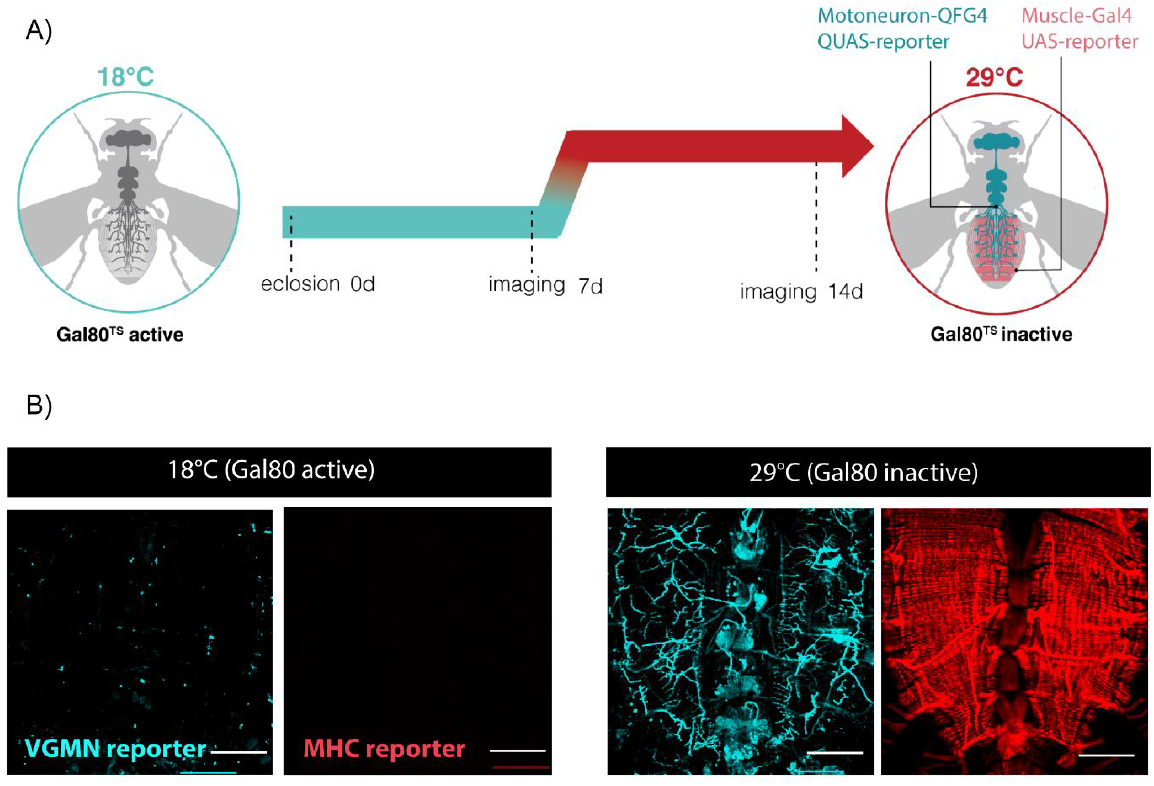
Gal4 and QFG4 can be simultaneously regulated by temperature sensitive Gal80. A) Schematic of QFG4 and Gal4 activators in presence of tsGal80 at 18°C. tsGal80 is active and can bind to the activation domain of both QFG4 and Gal4 simultaneously, hence the absence of expression from the QUAS- and UAS-reporters. When flies are shifted to 29°C, tsGal80 becomes inactive and QUAS- and UAS-reporters can be expressed. B) Left panel: Representative image of 7-days-old adult abdominal muscle field, in which tsGal80 is suppressing both VGMN-QFG4 and MHC-Gal4 drivers, when maintained at 18°C. Punctae visible in the 18°C condition are secondary antibody aggregates. Right panel: Representative image of 14 days old flies transferred at 29°C and kept for 7 additional days. tsGal80 is inactive and VGMN-QFG4 and MHC-Gal4 are simultaneously driving expression in otor neurons and muscles of UAS-CD8::GFP (red) and QUAS-mCherry (blue). Scale bar : 100 µM

## Discussion

Progressive advancements, innovations and refinements of binary gene expression systems have been driven by the demand for increasingly sophisticated genetic manipulations in *Drosophila* (2, 8, 24, 28, 82). However, exploiting the full arsenal of these tools in practice can be difficult and time-consuming. Vectors to utilise each binary system (Gal4/UAS, LexA/LexOp, QF/QUAS) were built in piecemeal manner (2, 24, 28) requiring bespoke strategies to introduce enhancer elements or transgenes into each system. The ability to clone and choose the genomic location of binary system elements is particularly important given that most large libraries of current binary genetic tools are located at just a few loci of favoured phiC31 ‘landing’ sites (41). In addition, temporal regulation of binary gene expression is often also critical, particularly where experiments must be restricted to a particular period, such as adulthood, without disrupting other periods, such as development for example. But again, the challenge of combining two temporal regulation components with two binary gene expression systems requiring a minimum of 6 genetic elements (i.e. Gal4 + UAS + tsGal80 + QF2 + QUAS + QS) plus the burden of controlling temperature (for tsGal80) and drug delivery (for QS) make constructing and then using such lines extremely challenging. Here, we have developed tools to address some of these challenges. Our modular vectors enable easy parallel cloning of transgenes or enhancer elements into similar vectors for all three binary genetic systems currently employed in *Drosophila* reducing the burden of molecular transgene construction and subsequent site-to-site genomic location expression variation. Second, we developed a new QF/Gal4 hybrid, QFG4, that can be regulated by Gal80, including temperature sensitive Gal80, reducing the assembly difficulty and experimental complexity of simultaneous negative regulation of two independent binary gene expression systems. Other QF and Gal4 hybrid molecules have previously been reported. A ‘split’ line using the QF DNA-binding domain with a separate compatible Gal4 activation domain has previously been demonstrated to be functional in *Drosophila* (31). However ‘split’ lines - even between two elements derived from Gal4 - are not regulable by Gal80 (85), without additional modifications (86). A contiguous QF DBD and Gal4 AD fused construct has been shown to be functional in *Zebrafish* (87). However, the *zebrafish* QFG4 variant is lacking the Gal4 middle domain and therefore, may be insensitive to Gal80 inhibition, a property retained in our novel hybrid molecule.

The modular vector series described here, pBID2, are designed to improve upon our previous pBID (38) UAS vector series. The original pBID UAS vector was designed to address the issues of phiC31 site-to-site transgene variability of expression by including gypsy insulator elements flanking the transgene (47) and unwanted Gal4 independent transgene ‘leak’ by utilising the DSCP (40) promoter element. While the pBID design successfully addressed these problems (38), it did come at a cost, notably that expression levels from this vector are reduced compared to hsp70 basal promoter based vectors (2, 88). In the pBID2 vector design, we have sought to overcome this limitation by exploiting a number of strategies including changing the 3’UTR sequence from hsp70 to the more potent p10 3’UTR (45) and utilising a Syn21 translational enhancer element (45), both intended to increase transgene expression. This approach enabled a *∼*150% increase in expression of transgenes from pBID2_UAS compared with the original pBID UAS vector, while still retaining the desirable feature of transgene insulation. On this improved backbone, we have also built pBID2_ LexOp and QUAS vectors, to complement our pBID2_UAS vector, enabling rapid parallel construction of transgenes suitable for all three binary systems. Our preferred method for introducing sequences into these vectors is Gateway cloning (44) and indeed all the transgenes constructed in this study were built using this method. While not aided by the (in our opinion) overly complicated product documentation, in practice Gateway cloning is an easy two step procedure, generation of an ‘entry’ clone, most often by topo cloning of a PCR product and then transfer to ‘destination’ vector, such as all the Gateway compatible pBID2 vectors described here, using a simple one-step reaction (Figure S1). However, as commercial gene synthesis costs continue to decline, synthesis of sequences directly into a Gateway compatible vector is increasingly becoming a simple and affordable solution to generate entry vectors with additional advantages such as the ability to exploit codon usage optimization. As a service to the *Drosophila* community, we have also built versions of the pBID2 vector series with multiple cloning sites designed to be backwards compatible with older vectors such as pUAST (2), for those who prefer restriction enzyme-based cloning methods (Figure S2).

We also evaluated the ability to produce multiple peptides from our vectors using ribosomal skipping sequences (62, 89). Very surprisingly, transgenes using the P2A ribosomal skipping sequence (63, 64), displayed incomplete skipping in motor neurons and produced both the expected multiple peptides but also likely ‘read-through’ larger unwanted fusion peptides. In contrast, the T2A ribosomal skipping sequence (62, 90) appeared completely efficient in these cells, producing only the expected peptides, from an otherwise identical construct. The P2A sequence ribosomal skipping sequence has been claimed to be equivalent to and even more efficient than the T2A sequence and is widely employed in cell lines (61), mammalian transgenes (91) and also some *Drosophila* constructs (62). Our results would caution the use of P2A to produce multiple peptides from a single open reading frame, at least for experiments designed for neuronal expression in *Drosophila*, and suggest the use of T2A ribosomal skipping sequences instead.

In addition to UAS, LexOp and QUAS pBID2 vectors, we also constructed pBID2 vectors that enable Gal4, LexA, QF2 and QFG4 expression under the control of enhancer sequences. Similar to the ‘responder’ vectors described above, these vectors also include flanking gypsy insulator sequences and utilise a combination of DSCP, Syn21 and p10 3’UTR sequences to optimise expression. We chose to utilise improved variants of each transcription factor designed to overcome prior limitations including the Gal4 4.2 variant designed to reduce Gal4-associated toxicity while maintaining high transcriptional activity (41), a hybrid LexA consisting of the LexA[DBD] and the QF[AD] designed to increase expression levels of LexOp transgenes and the QF2 variant of QF designed to reduce QF toxicity (27). To evaluate these vectors, we identified an enhancer element of vGlut, we dubbed VGMN, that enables expression of transgenes in motor neurons. VGMN regulated patterns of expression from pBID2 vectors were similar in all four binary systems, independent of landing site location, consistent with successful insulation of enhancer regulated transgenes. Interestingly however, comparison of levels of QF2 expression between Gal4, QF2 and QFG4 pBID2 lines revealed orders of magnitude more transgene expression from QF2 lines. While this may seem at first glance desirable, aberrantly high levels of exogenous gene expression could be a potential source of toxicity. Indeed we did note aggregation of some fluorescent protein reporters in QF2 lines, presumably due to very high levels of expression (27), that was not observed in Gal4, LexA or QFG4 lines. QFG4 expression levels were similar to Gal4 expression levels in our observations though the number of lines we have currently generated is limited. Consistently, QF/Gal4 hybrid constructs have previously been tested in *Zebrafish* where they also reduced transgene toxicity (87). However, high levels of QF DBD expression, as in QFG4, may still be more toxic than the GAL4 DBD (27). Consistent with this, we have anecdotally observed difficulty in generating QFG4 lines using enhancer sequences that drive very high levels of transgene expression, such as the MHC promoter (84). Nonetheless, given continuing issues of low transgene expression levels in LexA lines, QFG4 may be a generally useful second binary system of choice, with transgene expression levels more similar to those of Gal4.

Our primary motivation to generate QFG4 was to simplify the coordinate control of two binary gene expression systems by generating a QF variant that could be inhibited by Gal80. In our observations, QFG4 is effectively inhibited by both standard Gal80 and temperature-sensitive Gal80, with inhibition levels approaching those observed for Gal4. The strategy of adding the Gal4 activation domain to another DNA binding domain to enable Gal80 regulation has previously been demonstrated successfully for LexA[DBD]/Gal4[AD] hybrids (24), however the low expression level of LexOp transgenes by these lines has subsequently required the addition of more potent activation domains, such as the QF activation domain (27), as we have used our pBID2 LexQF constructs, to ensure adequate levels of LexOp transgene expression, thus losing the ability to be regulated by Gal80. The facility to regulate QFG4 by Gal80 has a number of advantages. First, QFG4 patterns of expression can be refined by existing large libraries of Gal80 lines (41), including mostly recently novel ‘trojan’ Gal80 lines (92). In contrast only a very limited number of QS lines currently exist. Second, as we have shown, QFG4 can be regulated by temperature sensitive Gal80 (93), simplifying temporal control, especially in combination with simultaneous Gal4/UAS expression, as opposed to requiring concurrent QS regulation by quinic acid administration. Drug regulable versions of Gal80 (14–18) should also be fully compatible with QFG4 lines, if current toxicity issues can be resolved (94). Temporal control of binary gene expression is a critical tool for a host of investigations, for example to discriminate effects of genes during development from effects during adulthood (14, 93). QFG4 lines should enable investigations where gene expression can be induced in discrete tissues simultaneously, to probe interactions between tissues or between receptors and ligands for example, using tissue specific Gal4 and QFG4 constructs. Constructing such lines, while still not trivial, is somewhat facilitated by the reduction in the number of components required. Moreover, all pBID2 constructs, including QFG4 constructs, use mini-white to allow selection (2, 95, 96) of transgenic animals which can ease the generation of complicated recombinant lines. When the mini-white marker gene is no longer needed, it can be removed, for example, using the “white eraser” method (97). In this study, we generated lines which allowed temperature sensitive Gal80 regulated expression of QUAS transgenes in motor neurons and UAS transgenes in muscles, enabling the coordinate and simultaneous manipulation of pre and postsynaptic partner cells. We envision the pBID2 tool set and QFG4 reagents will enable many other novel types of manipulations that were until now technically hindered, to probe molecular, cell and tissue interactions with temporal precision.

## Limitations of the Study

This study only examined transgene expression in *Drosophila melanogaster* and not other species. Comparison of Gal4, QF2 and QFG4 expression levels was only performed in lines employing the VGMN enhancer element and a limited number of QFG4 lines have currently been generated.

## Acknowledgments

We are grateful to Steven Stowers and Christopher G. Potter for reagents. We would like to thank members of the McCabe laboratory: Emma Källstig, Medha Raman, Samuel Vernon, Rebecca Smith and Chiara Paolantoni for critical reading of the manuscript. This work was supported by the Swiss National Science Foundation grant numbers: 31003A_179587 and 320030-232324 to B.M.

## Author Contributions

Conceptualization, B.M.; Investigation, B.M., E.R., Writing – Original Draft, Review & Editing, Funding acquisition, B.M.

## Materials and Methods

### Plasmid construction

The entire pBID2 vector series have been deposited at Addgene, a non-profit plasmid repository //www.addgene.org/Brian_McCabe/.

### pBID2 Gateway activator vector series

A three-fragment assembly strategy was used to generate the pBID2 activator vectors. Unique 8-cutter restriction sites were introduced between fragments to allow flexible insertions or exchanges. The following DNA fragments were amplified by PCR using Platinum SuperFi polymerase (Invitrogen): (1) The Gateway cassette–DSCP–IVS fragment was amplified from pBID1-G (Addgene # 35195) and flanked by PacI and SbfI sites. (2) Gal4, LexQF, or QF2 coding sequences were amplified from pBID1_G-Gal4.2, pCasper_act5bc-LexQF (Addgene #61310), and pQF2WB (Addgene # 61312), respectively, and flanked by PmeI and SbfI sites. (3) The p10 3^*′*^UTR was amplified from pJFRC81_10xUAS-IVS-Syn21-GFP-p10 (Addgene # 36432) and flanked by SbfI and XbaI sites. The three PCR fragments were assembled using HiFi DNA Assembly (Invitrogen). Each assembled construct was digested with PacI and XbaI and ligated into the linearized pBID1 backbone (Addgene # 35190), yielding in **pBID2_G-DSCP-Gal4-p10, pBID2_G-DSCP-LexQF-p10**, and **pBID2_G-DSCP-QF2-p10**. To generate pBID2_G-DSCP-QFG4-p10, the Gal4 DNA-binding domain (amino acids 1–147) was replaced with the QF2 DNA-binding domain (amino acids 1–183) using HiFi DNA Assembly. All vectors were transformed and amplified in ccdB-resistant competent cells (Invitrogen).

### pBID2 Gateway responder vector series

For the responder vectors, pBID2_20xUAS-G, pBID2_13xLexOp-G, and pBID2_10xQUAS-G were generated as follows. **pBID2_20xUAS-G**: The 20xUAS fragment was isolated from pBID_20xUAS-Hsp70-G-SV40 (gift from S. Stowers) via HindIII/AatII digestion and was inserted in place of the 10xUAS sequence in pBID1_G, yielding pBID_20xUAS-DSCP-G-SV40. **pBID2_13xLexOp-G**: The 13xLexOp fragment was isolated from pBID_13xLexOp-Hsp70-G-SV40 (gift from S. Stowers) and was inserted in place of the 10xUAS element in pBID1_G through HindIII/AatII digestion–ligation, resulting in pBID_13xLexOp-DSCP-G-SV40. **pBID2_10xQUAS-G**: The 10xQUAS element was amplified from p10xQUAS-CsChrismson (Addgene #163629), flanked by HindIII/AatII sites, and inserted in place of the 10xUAS sequence of pBID1_UAS-G, yielding pBID_10xQUAS-DSCP-G-SV40. For each of these responder vectors, the p10 3^*′*^UTR was amplified from pJFRC81_10xUAS-IVS-Syn21-GFP-p10 (Addgene #36432) and flanked by XbaI and HpaI sites. The SV40 polyA sequence was replaced with the p10 3^*′*^UTR using XbaI/HpaI digestion–ligation, resulting in **pBID2_20xUAS-DSCP-G-p10, pBID2_13xLexOp-DSCP-G-p10**, and **pBID2_10xQUAS-DSCP-G-p10**. All vectors were transformed and amplified in ccdB-resistant competent cells (Invitrogen).

### pBID2 multiple cloning site activator and responder vectors

Site-directed mutagenesis was used to replace the Gateway cassette with a multiple cloning site (MCS) in each pBID2 vector. The vector backbone, excluding the Gateway cassette, was PCR-amplified using primers with half of the MCS sequences on the 5’ primer and the other half on the 3’ primer. The amplified products were re-ligated using the KLD kit (New England Biolabs), transformed, and amplified in TOP10 competent cells (Invitrogen).

### pBID2 activator constructs with enhancers

A two-step cloning strategy was used to generate activator constructs containing specific enhancers. The vGlut motor neuron enhancer (VGMN) (78) was PCR-amplified from Canton-S genomic DNA and cloned into pCR8/GW-TOPO (Invitrogen), generating pCR8/GW-VGMN. The VGMN sequence was then transferred into each of the four pBID2_G driver vectors using LR Clonase II (Invitrogen), resulting in **pBID2_VGMN-Gal4, pBID2_VGMN-LexQF, pBID2_VGMN-QF2, pBID2_VGMN-QFG4**. The Synaptobrevin enhancer (nSyb) (40) was PCR-amplified from pBS-KS+ *retro*-Tango (panneuronal) (98) plasmid and cloned into pCR8/GW-TOPO (Invitrogen), generating pCR8/GW-nSyb. The nSyb sequence was then transferred into pBID2_G-QFG4 vector using LR Clonase II (Invitrogen), resulting in **pBID2_nSyb-QFG4**.

### pBID2 Responder Constructs with reporters

For each responder construct, the coding sequence was amplified by PCR from the templates indicated below. A Syn21 translational enhancer sequence was added immediately upstream of the start codon in all constructs. **pBID2_UAS-dsEGFP**: The dsEGFP coding sequence was amplified from pd2EGFP-N1 (99). **pBID2_QUAS-GtACR1::EYFP**: The GtACR1::EYFP coding sequence was amplified from pJFRC7-GtACR1-EYFP (Mauss et al., 2017). **pBID2_20xUAS-H2A::GFP-T2A-mKO2::CAAX** and **pBID2_10xQUAS-H2A::GFP-T2A-mKO2::CAAX**: Both construct were assembled from three PCR fragments using HiFi DNA Assembly (Invitrogen): (1) The Histone 2A (H2A) coding sequence was amplified from pBID LexOp-H2A-mCherry (gift from Prof. Steven Stowers). (2) The sfGFP sequence was amplified from pHD-sfGFP Scareless dsRed (Addgene #80811). (3) The mKO2 sequence was amplified from pCS2+ ChMermaid S188 (Addgene #53617). A CAAX membrane-targeting motif (54, 100) was added downstream of mKO2 using the reverse primer. The ribosomal skipping sequence T2A (Diao et al., 2015) was inserted between sfGFP and mKO2 using overlapping primers for sfGFP (reverse) and mKO2 (forward). **pBID2_13xLexOp-H2A::RFP-T2A-TFP::CAAX**: This construct was generated using a similar three-fragment HiFi assembly strategy: (1) The H2A sequence was amplified from pBID2 LexOp-H2A::mCherry (gift from Prof. Steven Stowers). (2) The RFP sequence was amplified from pBI-G-DSCP_RFP-myc-SV40 (McCabe Lab). (3) The TFP sequence was amplified from pJFRC_MUH-dBrainbow (Addgene #29327). A CAAX membrane motif was added downstream of TFP using the reverse primer. The T2A ribosomal skipping sequence was inserted between RFP and TFP using overlapping RFP reverse and TFP forward primers. The full-length coding sequences of all reporter constructs were cloned into pCR8/GW-TOPO (Invitrogen) and subsequently transferred into the appropriate pBID2_enhancer-G destination vector using the LR Clonase II kit (Invitrogen). **pBID2_20xUAS-H2A::GFP-P2A-mKO2::CAAX**: This construct was generated from **pBID2_20xUAS>H2A::GFP-T2A-mKO2::CAAX** by replacing the T2A sequence with a P2A sequence using PCR-based site-directed mutagenesis (KLD kit, New England Biolabs).

All constructs were sequence-verified by Sanger sequencing (Microsynth). Primer sequences are provided in Appendix 1 – Key resources table.

### Transgenic Drosophila lines generation

Transgenes were inserted into the following landing site using established methods. (Genetivision, Stafford, USA). pBID2_UAS-dsEGFP was inserted in attP40 (chromosome II), QUAS-GtACR1::EYFP was inserted in attP2 (chromosome III), VGMN-QF2, VGMN-Gal4, VGMN-LexQF, and VGMN-QFG4 were inserted in JK22C (chromosome II). LexOp-H2A::RFP-T2A-TFP::CAAX, UAS-H2A::GFP-T2A-mKO2::CAAX, UAS-H2A::GFP-P2A-mKO2::CAAX, VGMN-LexQF, VGMN-Gal4 and VGMN-QF2 were inserted in JK66B (chromosome III). QUAS-H2A::GFP-T2A-mKO2::CAAX, VGMN-QFG4 and nSyb-QFG4 were inserted in VK0005 (chromosome III).

### *Drosophila* stocks

A full list of the stocks employed is in Appendix 1 – Key resources table. All lines were raised on standard media at 25°C, 50% room humidity.

### Temporal control of transgene expression in larvae and adults

The Gal4/tsGal80 and QFG4/tsGal80 systems were used to achieve temporal control of UAS- and QUAS-reporters expression in muscle and motor neurons during larval and adult stages. MHC-Gal4 was used to drive UAS-GFP in muscle and VGMN-QFG4 was used to drive QUAS-mCherry in motor neurons. For larval experiments, two separate crosses were established: one maintained at 18°C to allow tsGal80 mediated suppression of transgene expression, and another maintained at 29°C to promote tsGal80 degradation and transgene activation. Third instar larvae from each condition were collected and analysed to assess the effects of tsGal80-dependent regulation. For adult experiments, a single cross was maintained at 18°C to suppress transgene expression by tsGal80 until 7 days post-eclosion. Adult flies were then shifted to 29°C to inactivate tsGal80 and allow UAS/QUAS transgene expression. Flies were collected at day 7 just before the temperature shift and at day 14, 7 days after the temperature shift. Both time point were analysed to evaluate transgene expression and the effects of tsGal80 regulation. Motor neurons reporter signal of QUAS-mCherry was enhanced by a staining with anti-DsRed rabbit primary antibody and anti-rabbit-546 secondary antibody (Appendix 1 - Key resources table).

### Western blot analysis

Adult fly heads expressing transgenes under the control of GMR-Gal4 (101) were homogenized in lysis buffer containing 137 mM NaCl, 20 mM Tris·HCl (pH 7.5), 1% Nonidet P-40, 10% (v/v) glycerol, Complete Ultra protease inhibitor cocktail (Roche), and PhosSTOP phosphatase inhibitor (Roche). Homogenates were sonicated three times for 5 s on ice and centrifuged at 12,000 × g for 10 min at 4°C to remove debris. Supernatants were collected, and 30 µg of total protein per sample was mixed with loading buffer to yield final concentrations of 10% glycerol, 2% (w/v) SDS, 5% (v/v) β-Mercaptoethanol, and 0.01% bromophenol blue. Samples were denatured at 70 °C for 10 min, resolved on 4–12% Bis-Tris WedgeWell minigels (Invitrogen) using MES–SDS running buffer, and transferred to iBlot PVDF membranes (Invitrogen). Membranes were blocked for 1 h at room temperature in PBST (PBS containing 0.1% Triton X-100) supplemented with 5% (w/v) non-fat dry milk. Blots were incubated overnight at 4°C with mouse anti-GFP primary antibody (1:10,000; Origin) diluted in PBST with 5% milk. After washing in PBST, membranes were incubated for 1 h at room temperature with the corresponding horseradish peroxidase (HRP)-conjugated secondary antibody (1:20,000) diluted in PBST with 2.5% milk. Following extensive washes in PBST, immunoreactive bands were visualized using WesternBright Sirius chemiluminescent substrate (Advansta) and imaged with an Amersham Imager 680 system (GE Healthcare). To confirm equal protein loading, membranes were stripped using Restore Stripping Buffer (Pierce) for 10 min at room temperature, washed in PBST, and re-probed with mouse anti-α-Tubulin antibody (1:20,000; Sigma-Aldrich). Protein band intensities were quantified using ImageJ software, and GFP signals were normalized to their corresponding α-Tubulin levels. Data are expressed as a percentage of control values.

### *Drosophila* larva and adult tissue dissection and Immunostaining

Dissection and immunostaining of *Drosophila* third instar (L3) larvae were performed as previously described (102, 103). Adult abdominal wall dissections were carried out following established protocols (104). Following dissection, samples were fixed in 4% Paraformaldehyde for 20 minutes and subsequently washed several times in PBST (PBS containing 0.1% Triton X-100). When immunostaining was performed, the preparations were first blocked for 30 minutes in PBST supplemented with 5% normal goat serum (PBSTG). The samples were then incubated overnight at 4 °C with primary antibody diluted in PBSTG: rabbit anti-DsRed (1:1000; Takara). After incubation with primary antibodies, the samples were thoroughly washed in PBST and incubated for 2 hours at room temperature with the appropriate secondary antibodies and dyes. Secondary antibodies included goat anti-mouse IgG conjugated to Alexa Fluor 488 (1:500; Invitrogen), goat anti-rabbit IgG conjugated to Alexa Fluor 555 (1:500; Invitrogen). To visualize neuronal membranes, Alexa Fluor 647-conjugated anti-HRP (Jackson ImmunoResearch) was used. Muscle fibres were labelled with Alexa Fluor 647-conjugated phalloidin (1:500; Biotium). After staining, samples were mounted in Vectashield mounting medium (Vector Laboratories) and imaged using either a Zeiss LSM 700 upright confocal or an Olympus FV4000 inverted confocal microscope. All images were processed using Image J.

### Luciferase assay

20-30 adult fly heads were homogenized in 150µl Glo Lysis Buffer (Promega) and incubated for 20 minutes at room temperature. Cellular debris were removed by centrifugation at 20,000 × g for 15 minutes at 4°C. An aliquot of 20 µL of each clarified homogenate was mixed with an equal volume of Steady-Glo Luciferase Reagent (Promega) and incubated for 10 minutes at room temperature in the dark. Luminescence was measured using a Spark multimode plate reader (Tecan). In parallel, total protein concentration from each homogenate was determined using the BCA Protein Assay Kit (Pierce). Background luminescence was subtracted and luminescence values were normalized to protein concentration for each sample. Results are reported as normalized luminescence units.

### Optogenetic and electrophysiological recordings from larval motor neurons

VGMN-GAL4 or QFG4 driver flies were crossed with UAS-GtACR1 and QUAS-GtACR1 reporter lines respectively, in the presence or absence of Tubulin-Gal80, on standard food supplemented with 1 mM all-trans-retinal (ATR) and maintained at 25°C. Third instar (L3) larval central nervous systems (CNS) were dissected and mounted dorsal side up on a Sylgard184 dish (Sigma) containing ice-cold external voltage-clamp saline (135 mM NaCl, 5 mM KCl, 5 mM TES, 36 mM sucrose, 26 mM NaHCO3, 1 mM NaH2PO4, 2 mM CaCl2, and 4 mM MgCl2; pH 7.2). Recording pipettes were pulled from borosilicate glass capillaries (GC120T-10; Harvard Apparatus) to a tip diameter approximating that of a single nerve root and filled with external saline. Neuronal activity was recorded from a single ventral nerve cord (VNC) projecting nerve root. Signals were acquired in current-clamp mode using an Axon Multiclamp 700B amplifier (Molecular Devices), low-pass filtered at 1 kHz, digitized at 50 kHz using a Digidata interface (Molecular Devices), and recorded with pCLAMP software (Molecular Devices). Optogenetic stimulation was delivered using an OptoLED system (Cairn Instruments, Kent, UK) producing teal/blue light (515 nm) through the epifluorescence light path of a 40×/0.8 NA water-immersion objective (LUMPlan FI; Olympus). Emitted light was filtered using HQ535/30 (Chroma) or a standard GFP emission filter. Preparations were recorded for 6 minutes with alternating 2-minute light ON/OFF periods. Recording traces were analysed offline using Clampfit software, version 11 (Molecular Devices).

### Cell nuclei counting

2 colour-channels confocal images were acquired from L3 larval VNC with a 20x air objective. After 3D reconstruction, image were analysed with Vision4D 3.0.0 (Arivis AG) as previously described (66). For each channel and the overlapping signal, a Blob Finder algorithm (105) was applied to detect nucleus and segment bright rounded three-dimensional sphere-like structures in the images with 4.5 µm set as the diameter. Segmented objects with volume less than 15 µm3 were removed from analysis by segmentation filter to avoid unspecific signals. Subsequently, the number of nuclei, included in VNC segment A4-A6, were output from Vision4D.

### Statistical analysis and reproducibility

Statistical analyses were performed using GraphPad Prism version 9.0.1 (GraphPad Software). Western blot data were analysed using unpaired two-tailed t-tests, while luciferase assay data was analysed using one-way ANOVA followed by Tukey’s multiple comparisons tests. Sample sizes are indicated in the corresponding figure legends. Throughout the text, figure legends, and tables, n refers to the number of biologically independent samples.

## Appendix 1

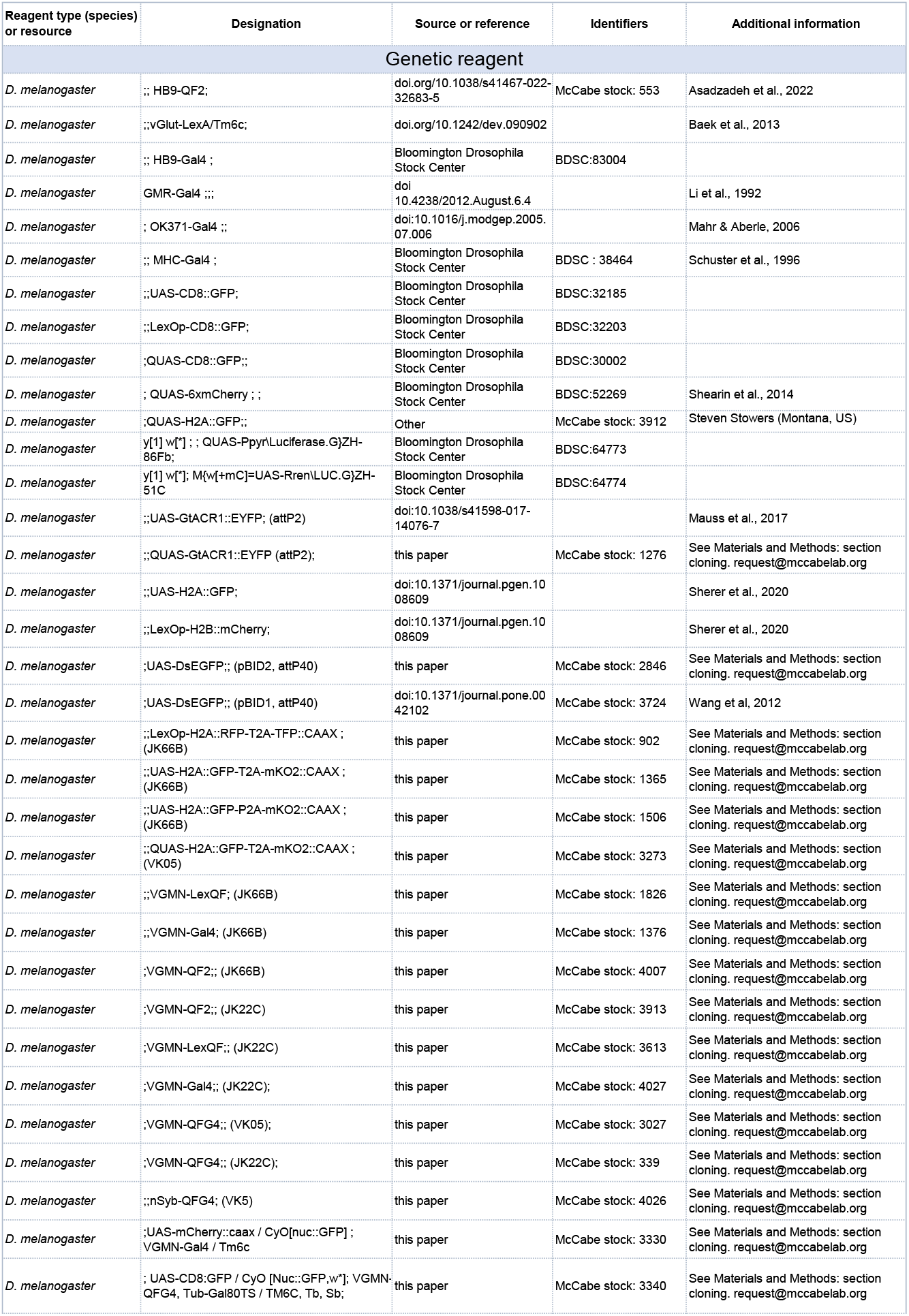
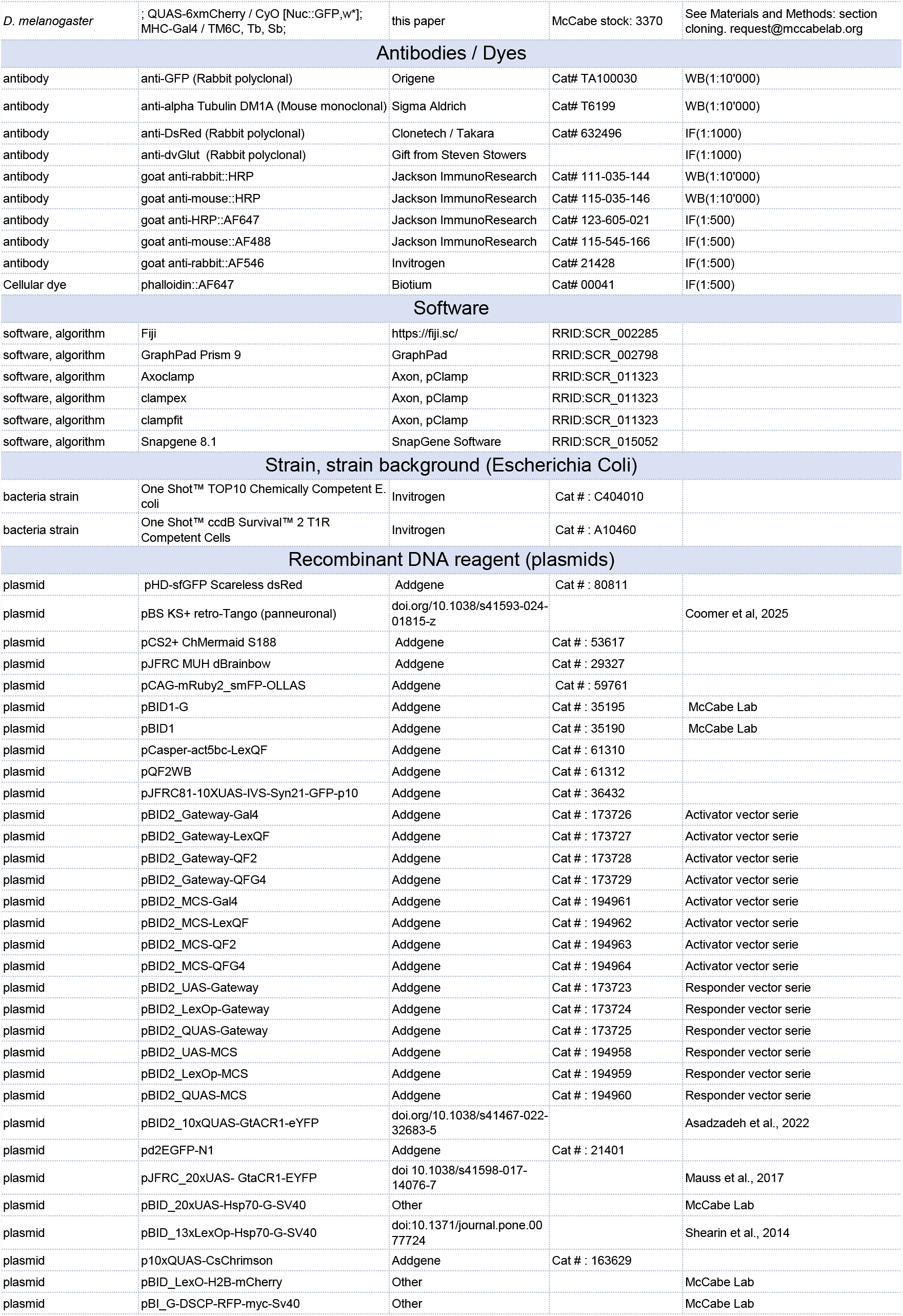
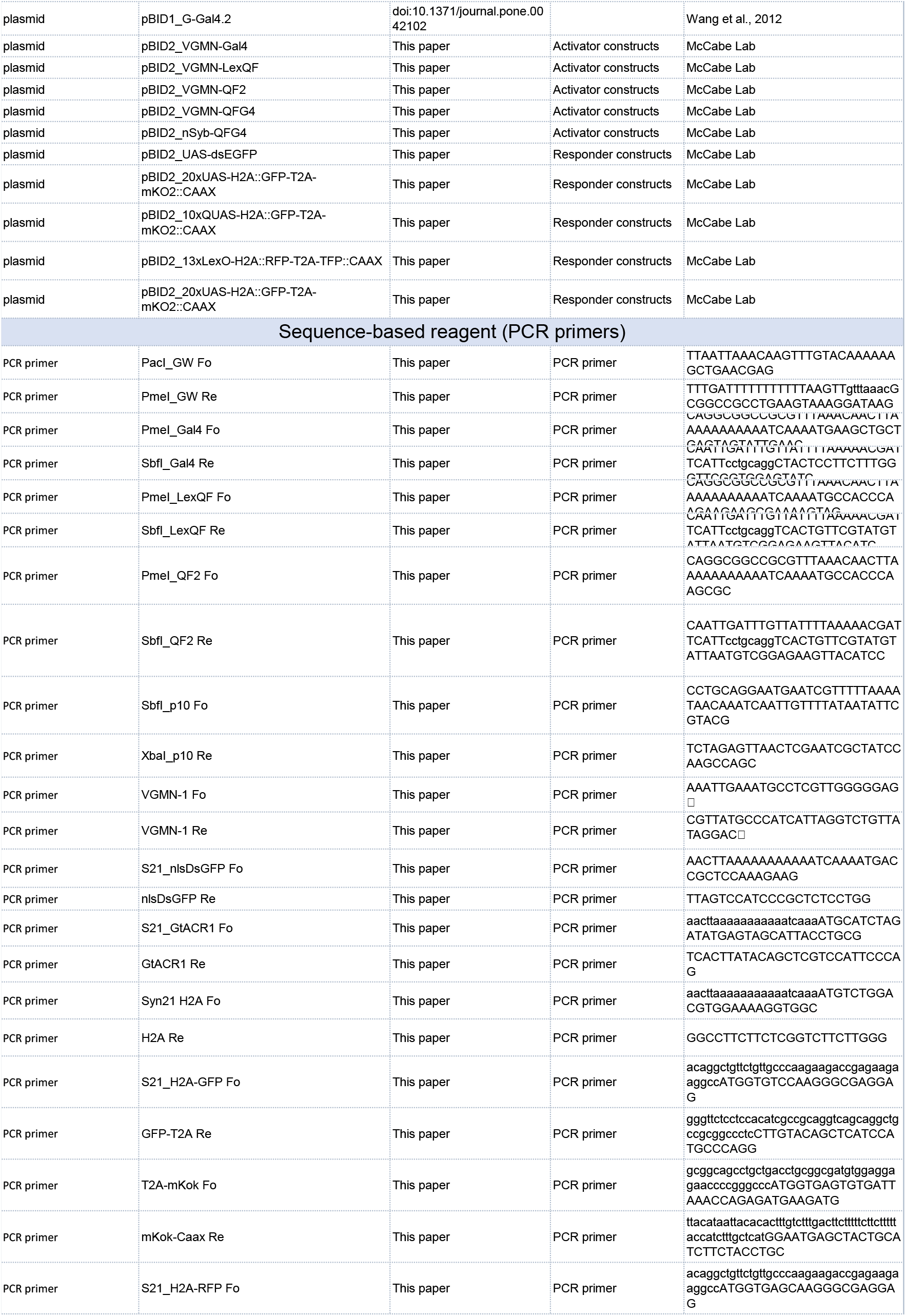
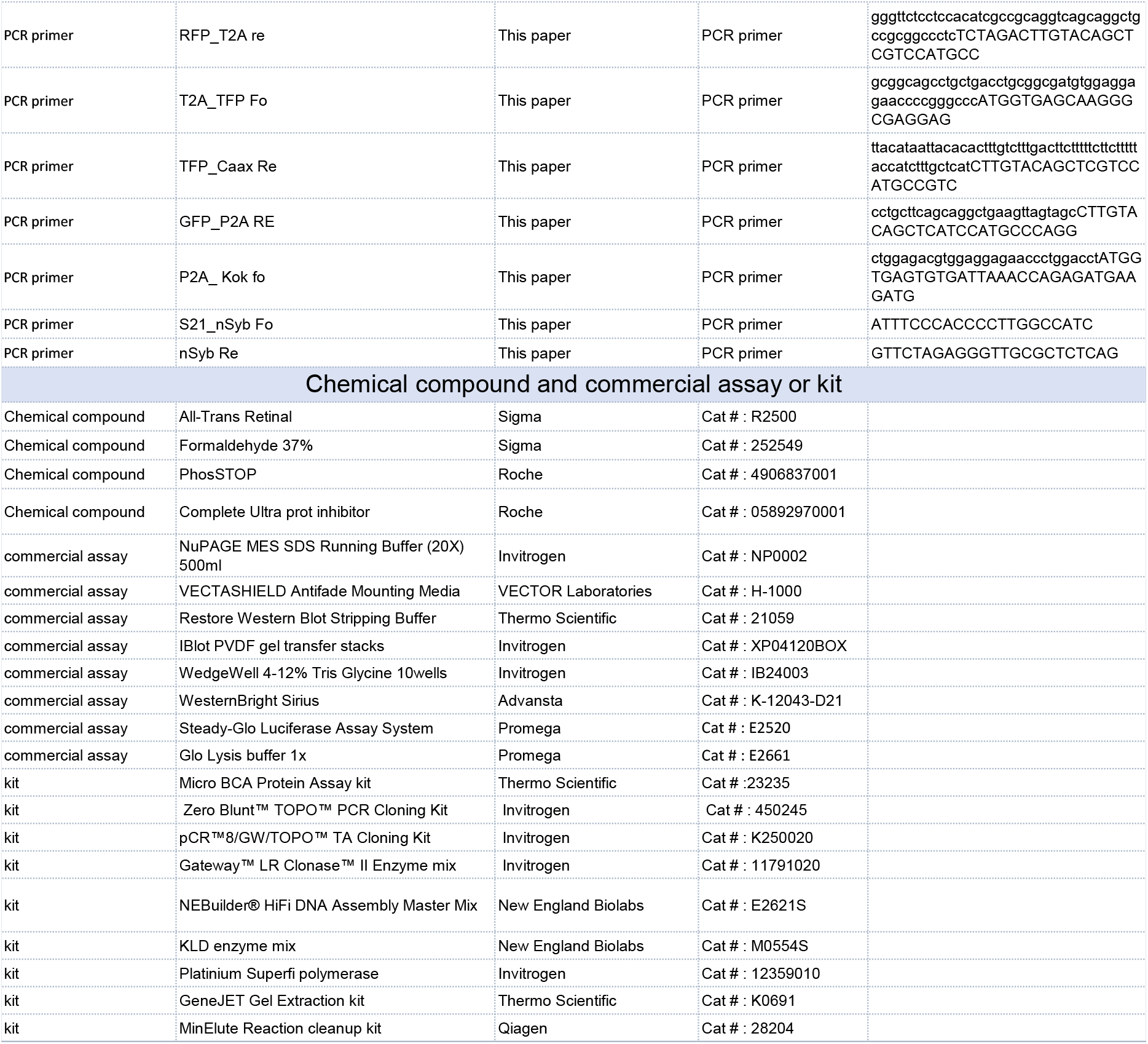

## Supporting information

**Figure S1.**
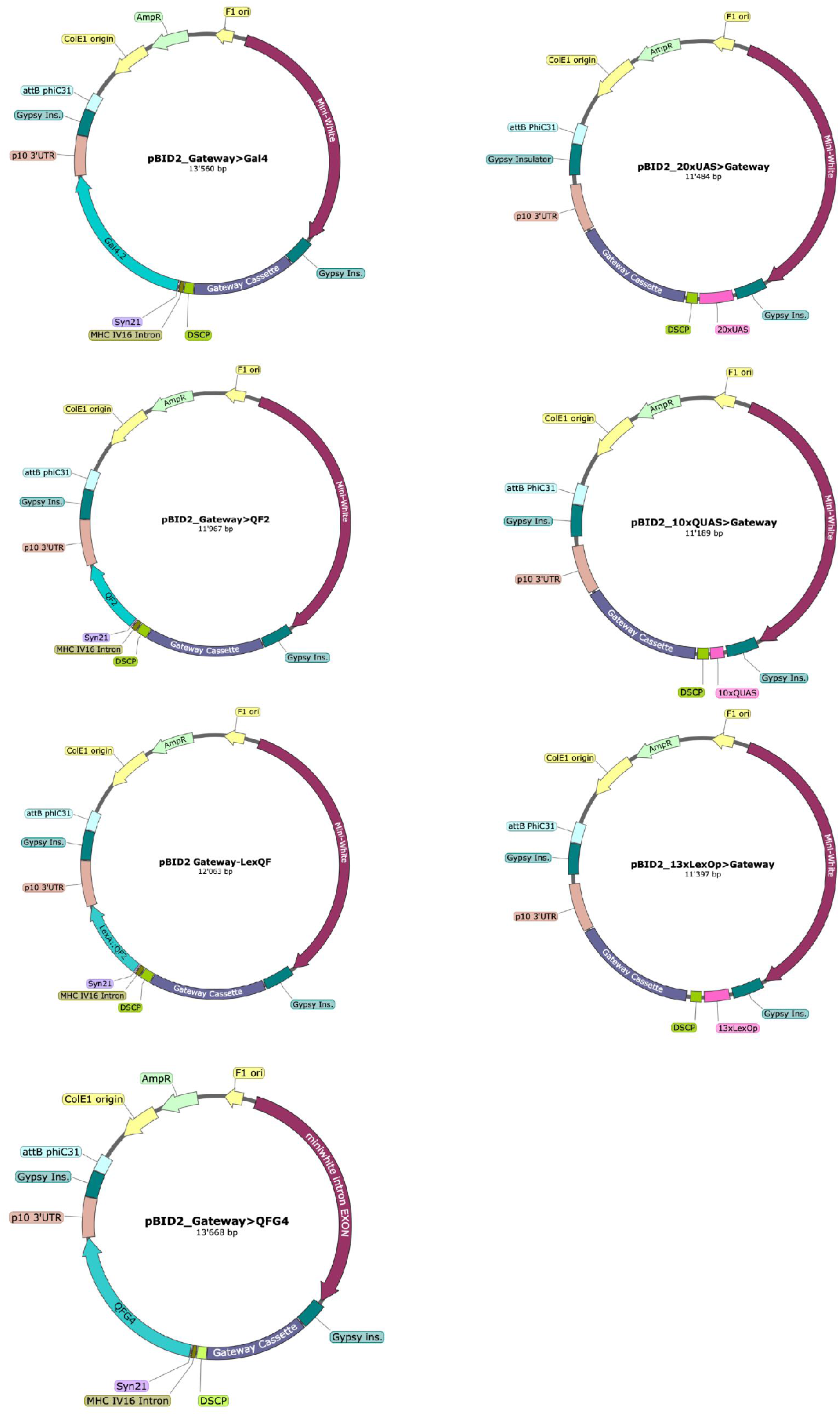
pBID2 activator and responder maps. Gateway version of the vectors.

**Figure S2.**
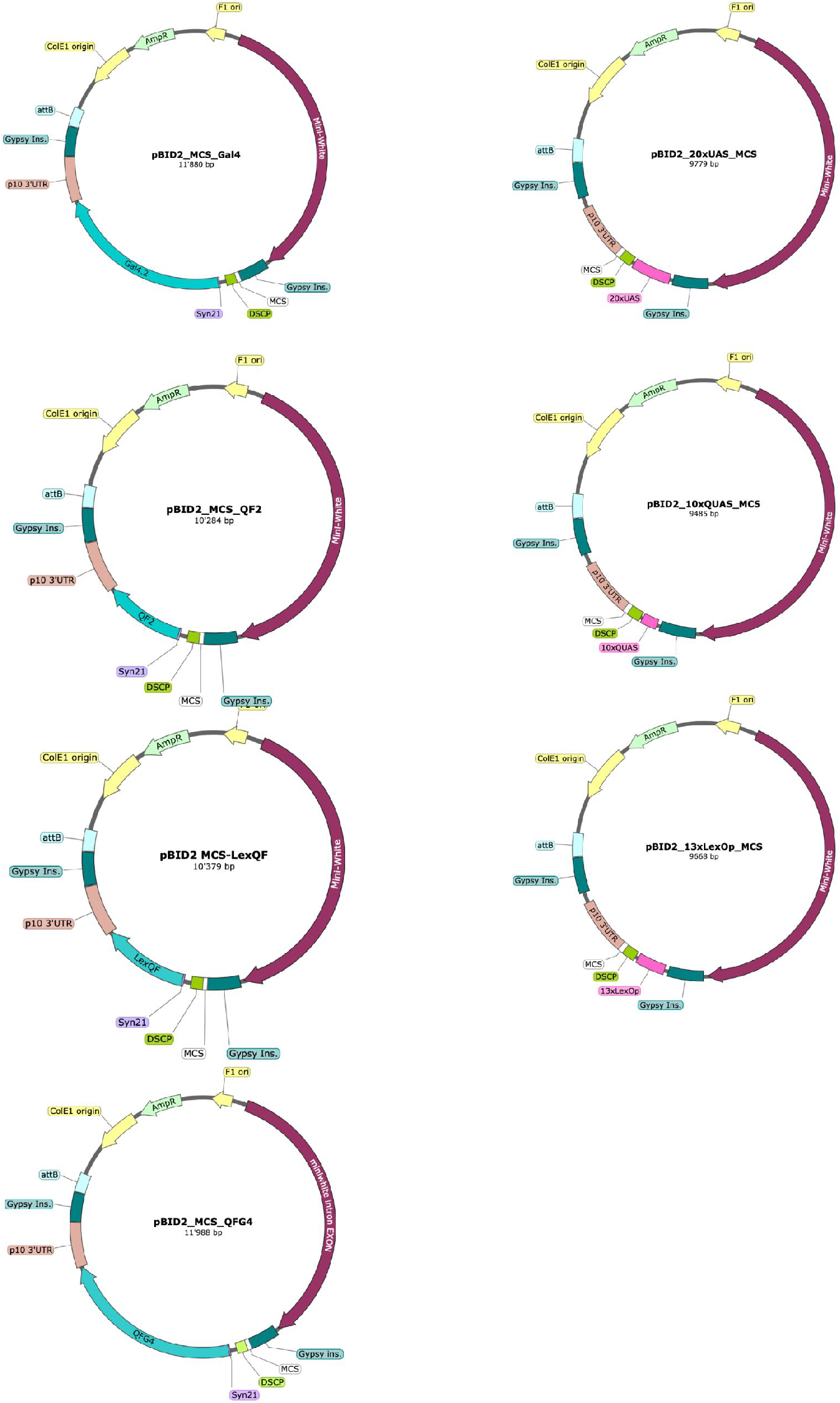
pBID2 activator and responder maps. Multiple cloning site version of the vectors.

**Figure S3.**
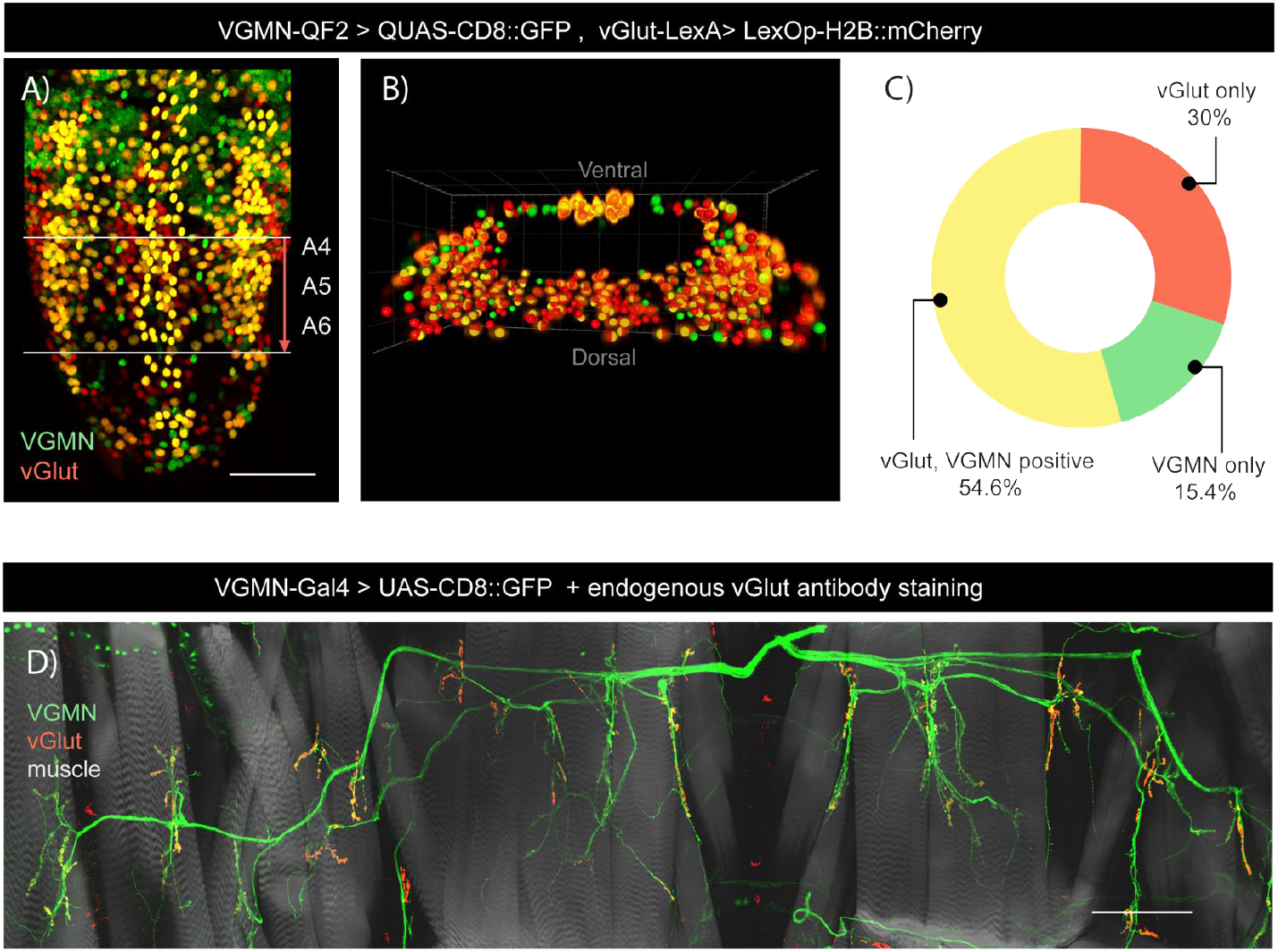
Number of VGMN labelled neurons vs all CNS glutamatergic neurons. A-C) Cell counting of vGlut neurons (OK371-Gal4) (red), VGMN neurons (VGMN-LexQF) (green) and neurons expressing both markers (yellow). A) confocal image of larval VNC showing segment A4-A6 in which cells were counted. B) 3D reconstruction of segment A4-A6 and quantification of cell nucleus with Arivis software. C) Cell number for VGMN+vGlut, vGlut only and VGMN only labelled nuclei. *n*=6. Scale bar: 50 µM D) Representative image of VGMN labelled motor neurons (in green) co-labelled for synaptic vGlut (in red). Scale bar 100 µM.

**Figure S4.**
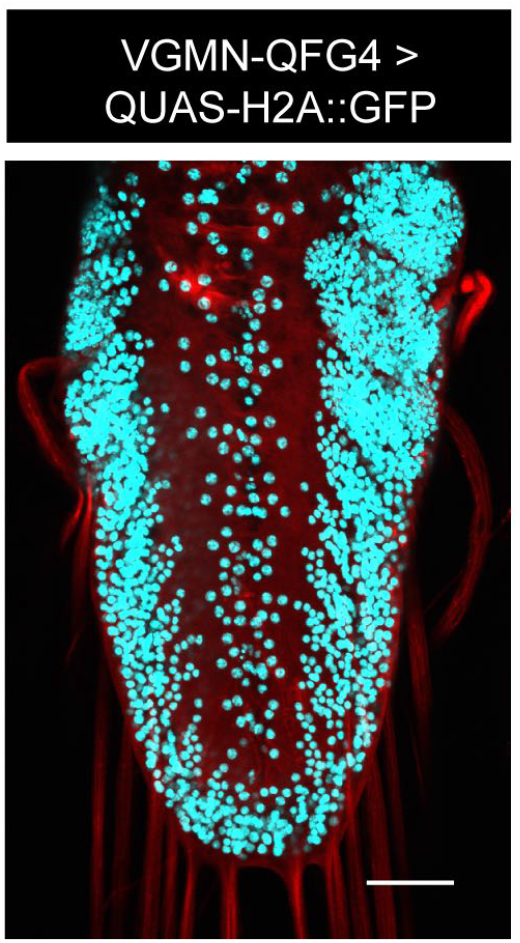
nSyb-QFG4 driver. Representative image of nSyb-QFG4 in the larval VNC (labelled with HRP-647 (red)) driving a QUAS-TFP nucleus reporter (blue). Scale bar 50 µM.

**Figure S5.**
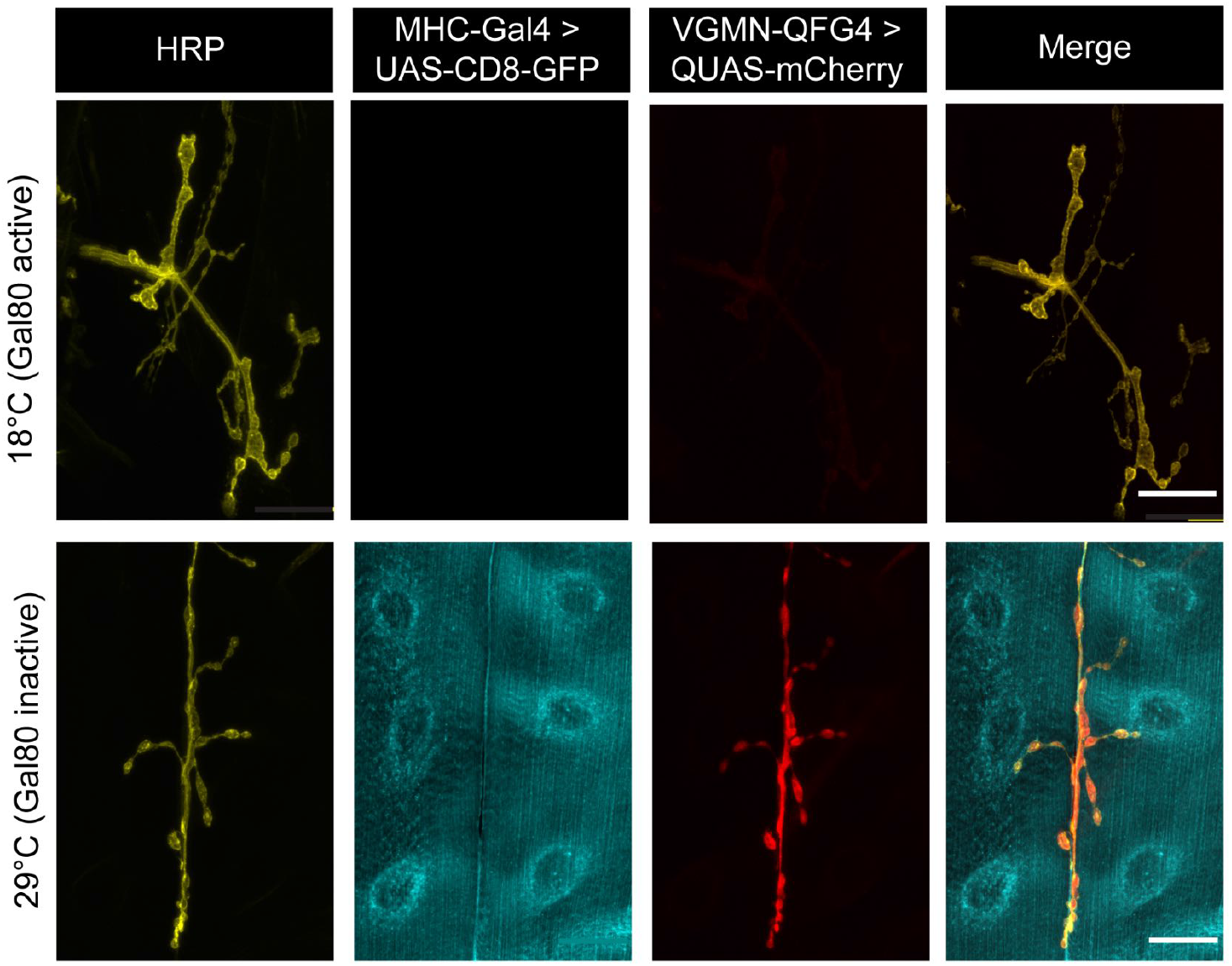
Simultaneous control of Gal4 and QFG4 by temperature sensitive tsGal80 in larva. Top row: Representative image of L3 larva raised at 18°C, in which tsGal80 is suppressing both reporter expression in nerve terminals (VGMN-QFG4) and in muscle (MHC-Gal4). Bottom row: Representative image of L3 larva raised at 29°C. tsGal80 is inactive and VGMN-QFG4 and MHC-Gal4 simultaneously can express both reporters, UAS-CD8:GFP (blue) and QUAS-mCherry (red). Left image: visualisation of neuronal terminals with HRP staining (yellow). Scale bars: 25 µM.

